# Microbiome-mediated resilience and cross-generational consequences in male *Drosophila* exposed to combined environmental stressors

**DOI:** 10.1101/2025.05.16.654542

**Authors:** Komal Maggu, Abhishek Meena, Alessio N. De Nardo, Benjamin Eggs, Sonja H. Sbilordo, Rawaa Al Toma Sho, Stefan Lüpold

**Affiliations:** Department of Evolutionary Biology and Environmental Studies, University of Zurich, Winterthurerstrasse 190, 8057 Zurich, Switzerland

**Keywords:** Microbiota, thermal fertility, pesticide exposure, inter-generational effects, fruit fly, stress resistance

## Abstract

Environmental stressors like heat extremes and pesticide exposure can significantly threaten insect reproduction, yet the interplay of these stressors and the potential mitigating role of the gut microbiome remain poorly understood, particularly across generations. This study investigated the interactive effects of acute heat stress and sublethal imidacloprid exposure on male *Drosophila melanogaster* reproductive success and the subsequent fitness of their offspring. We manipulated the gut microbiome of male flies (germ-free or colonized with one or five bacterial species) and subjected them to individual or combined stress. Our findings reveal that combined stress synergistically impairs male fitness traits, an effect partially buffered by higher microbiome diversity. We further demonstrate cross-generational consequences, with paternal stress exposure also impacting offspring fitness. Our results highlight the crucial role of the gut microbiome in mediating resilience to environmental stress and underscore the importance of considering multi-stressor and intergenerational effects in ecological risk assessments.

## Introduction

The increasing prevalence of human-induced stressors poses a significant challenge to global biodiversity (Navarro-Ortega *et al*. 2015; Piggott *et al*. 2015b, a; Przeslawski *et al*. 2015; Segner *et al*. 2014), with climate change and pesticide being key drivers of biodiversity loss and their interaction presenting unique ecological risks (Beketov *et al*. 2013; Heino *et al*. 2009; Stehle & Schulz 2015). Rising temperatures are projected to intensify pesticide use due to intensified pest outbreaks (Deutsch *et al*. 2018), while simultaneously altering the response of ectotherms to chemical exposure (Fournier-Level *et al*. 2016; Hooper *et al*. 2013; Noyes *et al*. 2009). For ectotherms, which lack internal temperature regulation (Deutsch *et al*. 2008; Paaĳmans *et al*. 2013), warming elevates metabolic rates and feeding activity (potentially increasing pesticide uptake), leading to energetic trade-offs between detoxification and essential functions like growth or reproduction (Jutfelt *et al*. 2021; Pörtner *et al*. 2017).

While the individual effects of temperature and pesticides are relatively well-documented (Dougherty *et al*. 2024; Wan *et al*. 2025), their combined impacts remain poorly resolved but have shown variable outcomes. Studies report additive (Arambourou & Stoks 2015), antagonistic (Meng *et al*. 2022), or synergistic (Wang et al., 2019) interactions, with synergism being most concerning for non-target species as their combined harm exceeds the sum of individual effects (Folt *et al*. 1999; Siviter *et al*. 2021; Verheyen & Stoks 2019, 2023). Such synergisms may arise from physiological trade-offs, such as heat stress diverting resources from detoxification (Dai *et al*. 2023) combined with a pesticide-induced antioxidant depletion, crucial for thermal resilience (Silva *et al*. 2025). This dual depletion could disproportionately harm gonads and gametes that are particularly prone to oxidative stress due to their high cell proliferation, metabolic rate and generation of reactive oxygen species (Simmons *et al*. 2018). By eroding reproductive resilience, synergisms threaten to accelerate biodiversity loss in ways that single stressors would not (Shahid *et al*. 2024).

Beyond direct physiological impacts, host-associated microbial communities (microbiomes) are increasingly recognized as mediators for environmental stress tolerance (Feldhaar 2011; Osborne *et al*. 2023; Sepulveda & Moeller 2020; Shokal *et al*. 2016). Commensal microbes can enhance host thermal performance (Renoz *et al*. 2019; Sepulveda & Moeller 2020), either as obligate or facultative symbionts (Dunbar *et al*. 2007; Gruntenko *et al*. 2017; Montllor *et al*. 2002; Russell & Moran 2006; Zhang *et al*. 2008), and a healthy microbiome can bolster insect resistance to insecticides through direct pesticide metabolism or by inducing the expression of host detoxification genes (Cheng *et al*. 2017; Jones *et al*. 2015; Kikuchi *et al*. 2012; Pang *et al*. 2018). Despite progress in studying individual stressors, how microbiomes modulate *combined* effects remains largely unknown, but microbiome disruption by one stressor could amplify susceptibility to another, potentially explaining observed synergisms (Karimzadeh *et al*. 2020; Mao *et al*. 2019; Zhang *et al*. 2008; Zhang, Cai, *et al*. 2021).

Another critical knowledge gap concerns stress effects on male reproduction. While much research focuses on acute toxicity or sterility thresholds (De França *et al*. 2017; Desneux *et al*. 2007; Walsh *et al*. 2014; Zwoinska *et al*. 2020), sublethal exposures can have significant fitness consequences. Sublethal effects on female reproductive function are relatively well-characterized, the studies on males are lagging behind but have already revealed that male fertility can be more vulnerable than female fecundity, at least to heat stress (Iossa 2019; Sales *et al*. 2018; Zwoinska *et al*. 2020). Heat and pesticides can impair spermatogenesis, sperm DNA integrity, and overall ejaculate quality (Aulsebrook *et al*. 2020; Dougherty *et al*. 2024; Walsh *et al*. 2019), key determinants fertlization success, particularly under competitive conditions in promiscuous species (Fitzpatrick & Lüpold 2014; Parker & Pizzari 2010; Pizzari & Parker 2009; Simmons & Fitzpatrick 2012).

Sperm damage can also extend the environmental stress effects across generations through epigenetic modifications or sperm-borne small RNAs (Bonduriansky *et al*. 2012; Morgan & Watkins 2019), affecting population stability and evolution (Aulsebrook *et al*. 2020; Crean & Immler 2021). While parental exposure to combined thermal and chemical stress has been shown to reduce offspring performance in some species (DeCourten & Brander 2017; Fathy 2024), other studies find no clear effects (Massot *et al*. 2021; Meng *et al*. 2022) or even reduced offspring vulnerability (Tran *et al*. 2018), indicating interspecific variation due to stressor intensity (including the thermal sensitivity of pesticides themselves), species-specific acclimation, or compensatory maternal provisioning. Understanding these cross-generational impacts is crucial for predicting long-term ecological consequences and evolutionary adaptation potential.

To address these knowledge gaps, we investigated the interactive effects of acute heat stress and sublethal pesticide exposure on the reproductive fitness of male *D. melanogaster*, focusing on microbiome mediation and cross-generational consequences. We combined sublethal acute heat stress (4 hours at 36°C), previously shown to negatively affect testicular function and the fertilizing capacity of sperm (Meena *et al*. 2024a, b), with exposure to imidacloprid, a widely used systemic insecticide targeting nicotinic acetylcholine receptors (nAChRs) in the central nervous system (Ihara *et al*. 2017). Sublethal imidacloprid exposure is known to affect various aspects of insect physiology and fitness (De França *et al*. 2017; He *et al*. 2011; Kang *et al*. 2018; Li *et al*. 2018; Xiao *et al*. 2016; Zhang *et al*. 2021b; Zhou *et al*. 2017). To isolate the role of the microbiome, we compared germ-free (axenic) males with males associated with defined bacterial communities: mono-association of *Acetobacter tropicalis* or *Lactobacillus brevis*, or a five-species consortium of *Acetobacter* and *Lactobacillus* species that are common in *Drosophila* (Broderick & Lemaitre 2012; Gould *et al*. 2018; Newell & Douglas 2014; Ridley *et al*. 2013). This gnotobiotic approach permitted precise assessment of microbial contributions to stress tolerance. Furthermore, we assessed the cross-generational consequences of paternal stress exposure on offspring fitness, testing if the father’s microbiome status influenced these effects while minimizing bacterial transmission to offspring to focus on direct paternal effects (Marshall *et al*. 2010; Moore *et al*. 2019; Mousseau *et al*. 2009).

Based on the evidence outlined above, we tested the following five predictions: (1) Combined sublethal heat and imidacloprid exposure will synergistically reduce male reproductive fitness. (2) Gut microbes will partially mitigate the negative impacts of individual and combined stressors on male reproductive fitness compared to germ-free males. (3) Different bacterial compositions will vary in their protective capacity, with the complex community offering greater resilience than either mono-association. (4) Paternal exposure to individual and combined stressors will negatively impact offspring fitness (cross-generational effects), with potential synergistic amplification. (5) The paternal microbiome composition will influence the magnitude of these cross-generational effects.

## Materials and Methods

### Experimental populations

Throughout this study, we used an outbred *D. melanogaster* population (LH_m_ wild-type strain), maintained under constant conditions of 25°C, 60% humidity, and a 12h:12h light:dark cycle. First, we generated axenic (germ-free) focal flies by collecting eggs (within six hours of oviposition) from four large population containers supplied with Petri dishes containing grape-juice agar medium (300 mL water, 670 mL grape juice, 22 g agar, 20 g sugar, 19 g yeast, 20 mL nipagin per litre) supplemented with a smear of yeast paste. Eggs were washed off the plates into a strainer, pooled, and dechorionated by immersion in 3.5% bleach for 2 minutes, followed by a rinse in 70% ethanol and three rinses in sterile water.

Next, eggs were aseptically transferred to sterile glass vials containing autoclaved standard fly food (75 g glucose, 100 g fresh yeast, 55 g corn, 10 g flour, and 15 mL nipagin antimicrobial agent per liter food media) supplemented with antibiotics [a cocktail of following antibiotics with their respective final concentrations: Ampicillin (50 μg/mL), Kanamycin (50 μg/mL), Erythromycin (10 μg/mL), Tetracycline (10 μg/mL)]. Vials were then sealed with sterile plugs and incubated under standard laboratory conditions until adult eclosion. All steps of vial preparation were conducted in a laminar flow chamber to avoid contamination.

Upon eclosion, axenic status was confirmed using two methods on a subsample of flies. First, homogenated flies were plated on Lysogeny Broth (LB) and de Man, Rogosa and Sharpe (MRS) agar plates, incubated, and checked for bacterial colony growth. Second, genomic DNA was extracted from pools of 3-5 putatively germ-free and conventional flies (positive control), respectively, using Quick-DNA midiprep plus kit following the instructions provided by the manufacturer. Universal primers (27F and 1492R) were used as forward and reverse primer, respectively. To prepare the PCR master mix, 10 μL of nuclease-free water, 0.5 μL of forward and reverse primer, 1 μL of template DNA and 13 μL Taq green master mix were mixed in a sterile eppendorf tube. PCR conditions were: initial denaturation at 95°C for 90s, followed by 35 cycles of denaturation at 95°C for 30s, annealing at 55°C for 30s and extension at 72°C for 90s, and final extension at 72°C for 5 minutes). PCR products were visualized on a 1% agarose gel under ultraviolet light to confirm the absence of bacterial DNA amplification in axenic samples and its presence in controls.

### Bacterial cultures and gnotobiotic associations

Axenic males were associated with defined compositions of microbial species, all originally isolated from the gut of wild-type flies (Obadia *et al*. 2017): *Acetobacter tropicalis* (*At*), and *Lactobacillus brevis* (*Lb*) for mono-associations based on their contrasting effects on male reproductive investments in our previous study (Maggu *et al*. 2025). For comparison, we also included axenic males and males inoculated with a defined five-species community consisting of common and abundant *Drosophila* gut bacteria (*A. tropicalis*, *A. orientalis, A. pasteurianus, L. plantarum, L. brevis*), resulting in four distinct microbial treatments.

Bacterial strains were cultured overnight in an incubator at 30°C: *Acetobacter* in MRS + mannitol medium with shaking, and *Lactobacillus* in MRS broth under stationary conditions. The following morning, cultures were checked for growth before measuring their optical densities (OD) at 600 nm and centrifuging at 7000 rpm for 5 minutes. Bacterial pellets were then resuspended in sterile phosphate-buffered saline (PBS) to a final concentration of 10^8^ cells/mL, following established protocols (Koyle *et al*. 2016; Newell & Douglas 2014).

To inoculate vials, 50µL of bacterial suspension (approx. 5×10^6^ CFUs) was added to sterile food vials, with equal representation of bacterial strains in the 5-species consortia. This bacterial density is comparable to previous reports for conventionally reared flies (Gould *et al*. 2018; Newell & Douglas 2014). Inoculated food vials were then allowed to air-dry in a laminar flow cell before introducing ten 1-day-old axenic males to each to generate gnotobiotic flies. Males were transferred to freshly inoculated vials with the same bacterial treatment every 48h for a total of four days to ensure stable colonization. Successful colonization with the intended bacteria was confirmed by plating homogenates of males from each treatment and checking for characteristic colony morphologies (see *Bacterial abundance of males*).

### Stress treatments

Following the 4-day colonization period, males were transferred to vials containing sterile food and exposed to pesticide and heat stress in a fully factorial design. Specifically, half of the males from each microbial group were subjected to food supplemented with imidacloprid, whilst the remaining half were maintained on uncontaminated standard food. We used a concentration of 1 ppm (parts per million) of imidacloprid as concentrations of 0.04 to 20 ppm have been used effectively in studies on larval or adult *D. melanogaster*, depending on the exposure regime (Charpentier *et al*. 2014; Frantzios *et al*. 2008; Martelli *et al*. 2020). These concentrations are low enough to avoid immediate lethality but sufficient to induce sublethal effects on behavior, reproduction, and physiology. Males were exposed to imidacloprid for an acute window of 18 h, following an additional thermal treatment of 4-hour in a hot water bath. To this end, half of the males from each microbiome and pesticide treatment were subjected to acute heat stress by transferring them to vials containing only sterile 1% agar (to prevent dehydration) and placing these vials in a water at 36°C for 4 hours. The remaining males (thermal controls) were treated identically but at benign 24.5°C. All males were then allowed to recover overnight in standard food vials under standard laboratory conditions.

### Hatching success and fertility

To quantify the fitness consequences of heat and pesticide exposure and their potential mitigation by the microbiome on the morning after the stress treatments, experimental males were individually mated to standard females of the same population. All females had been collected as virgins (within 7 hours of eclosion) and reared under axenic conditions along with the experimental males to minimize confounding contributions of female microbes to the measured outcomes. After mating ended, females were allowed to lay eggs for 24 h, and another 30 h later (allowing time for embryonic development), oviposition vials were frozen for later quantification of the proportion of eggs hatching as a proxy of fertilization success, and the total number of larvae estimating male-mediated fecundity. This assay was performed in two blocks, each with 25 males per treatment combination, totaling 800 experimental males (4 bacterial treatments × 2 pesticide treatments × 2 temperature treatments × 2 blocks × 25 males).

### Sperm competitiveness

To assess male sperm competitiveness, the above experimental design was repeated, but this time using standard females that had mated once with standard competitor males two days before the focal mating. These standard competitors were of the same strain but genetically modified to ubiquitously express green fluorescent protein (GFP), permitting paternity assignment against the focal wild-type males (Lüpold *et al*. 2012, 2020). Pairs were monitored for remating for up to 8 h. Remated females were then isolated in fresh vials for oviposition, while pairs not mating were separated and given another 8-hour mating opportunity on the following day. Females still not remating were excluded. Offspring paternity (focal vs. competitor) was assessed via GFP fluorescence to calculate P_2_ as the proportion of offspring sired by the second (focal) male. The treatment design of this assay was identical to the non-competitive experiment above, but this time with 30 trials per treatment combination in each of the two experimental blocks (i.e., 960 trials in total).

### Bacterial abundance in males

To investigate the impact of stress treatments on the bacterial load, colony forming units (CFUs) were compared between ten random males from each experimental treatment, surface-sterilized with 70% ethanol, and individually homogenized in 1.5 mL microcentrifuge tubes containing sterile PBS using a pestle. Homogenates were serially diluted to different dilutions at a final volume of 100 μL, then plated on MRS media for quantification of CFUs at appropriate dilutions (approx. 100 CFUs/plate). For the control groups with both *Acetobacter* and *Lactobacillus* species, MRS agar was supplemented with 10 μg/mL ampicillin to avoid the growth of *Lactobacillus,* whilst replicate plates were incubated in a CO_2_ atmosphere to exclude the growth of *Acetobacter*. The presence of only target bacterial species on plates was confirmed based on colony morphologies, with *A. tropicalis* producing small tan-brown colonies and *L. brevis* producing small yellowish colonies. CFUs per fly were calculated as: CFU = *C* × *D*/*P* × *V*, where *C* = number of colonies counted, *D* = dilution, *P* = volume plated, *V* = volume of fly homogenate (Koyle *et al*. 2016). For control males, with all the five bacterial species, bacterial colonies were identified by their morphology and total CFUs per fly was obtained by adding CFUs from all the five bacterial species.

### Cross-generational effects

To assess cross-generational impacts of paternal (F_0_) stress exposure, we quantified hatching success and fecundity in both male and female offspring (F_1_) of our experimental males. Within each experimental block, up to 20 freshly hatched larvae from each of ten randomly selected vials per parental treatment combination (i.e., 16 combinations) were transferred to rearing vials containing sterilized standard fly food. This immediate transfer minimized the potential acquisition of parental microbes via consumption of the egg chorion or contaminated food. These larvae developed under standard laboratory conditions.

Upon eclosion, 4-5 virgin F_1_ male and female offspring were collected from each rearing vial, separated by sex. At 2-3 days of age, three males and three females from each lineage were individually mated with a virgin standard fly of the opposite sex reared under axenic conditions (to avoid potential confounding effects of their microbiota). Following copulation, hatching success and fecundity were measured for these F_1_ flies as in their parents (see above). These offspring themselves were not subjected to any experimental treatment, thus assessing solely the cross-generational effects. This experiment was carried out in two independent blocks of 480 offspring per sex from 160 F_0_ males (i.e. total of 1,920 mating trials).

### Statistical analysis

All statistical analyses were performed in R version 4.4.3 (R Core Team 2025), using generalized linear models (GLMs) with a Poisson (fecundity) or binomial error distribution (hatching success, paternity shares) in the experimental generation, and the corresponding mixed-effects models (GLMMs) in the offspring generation with sire as a random effect to account for non-independence among siblings. Each model included microbiome composition (4-level factor) along with the binary thermal and pesticide treatments with all two- and three-way interactions as fixed effects, along with experimental block as a nuisance factor accounting for inter-block variation. Given that this factor only had two levels except for the assay of CFUs with three experimental blocks, thereby limiting estimates of random variance, we included it as a fixed effect in all models.

The binomial model of the relative abundance of Acetobacter versus Lactobacillus exhibited slight overdispersion (dispersion factor = 1.12), which we addressed by fitting a mixed-effects model with an observation-level random effect. In contrast, the remaining non-linear models showed moderate underdispersion (dispersion factors ≈ 0.9). Alternative model families such as Tweedie, negative binomial, and quasibinomial did not improve model fit, so the original models were retained for clarity and interpretability.

Model performance was checked using the performance package (Lüdecke *et al*. 2021), and output plots were generated was plotted using ggplot2 (Wickham 2016) based on means and 95% confidence intervals estimated via emmeans (Lenth 2025).

The impact of the different experimental treatments was also analyzed for the total bacterial abundances (CFUs) and, separately, for compositional shifts in the five-species consortia. For the former, we used a linear mixed-effects model with block as a random effect. For microbiome community composition in the five-species consortium, we quantified the species-specific abundances per sample, then applied a Hellinger transformation to these count data and computed Bray-Curtis distances using the *vegdist* function in the package vegan (Oksanen *et al*. 2025). We assessed treatment effects (temperature, pesticide, and their interaction) in a permutational multivariate analysis of variance (PERMANOVA) using *adonis2* in the same package. To visualize compositional shifts, we further ran non-metric multidimensional scaling (NMDS) (*metaMDS*) on the Bray-Curtis dissimilarities and fitted species vectors onto the NMDS ordination with *envfit*, which indicates the direction and strength of each species-specific association with the ordination space. We also compared the relative abundances at the genus level, using a binomial GLMM with block as a random effect and accounting for overdispersion by including an additional observation-level random effect.

## Results

### Hatching success

To assess the combined effects of heat stress, pesticide exposure, and microbiome composition on male reproductive function, we first investigated fertilization success, estimated as the proportion of eggs hatched out of all eggs laid by females. A binomial GLM revealed a significant three-way interaction between microbiome, temperature, and pesticide (*N* = 650, *χ*² = 15.19, *df* = 3, *p* = 0.002; Fig. 1A, Supplementary Table S1), highlighting that the combined effects of these stressors depended on the microbial composition of the focal males. Overall, males associated with all five bacterial strains exhibited the highest hatching success, followed by *At*- and *Lb*-associated males, with axenic males being most severely affected. These general patterns were also confirmed by the significant two-way interactions of temperature ’microbiome (*χ*² = 22.01, *df* = 3, *p* < 0.0001) and microbiome ’pesticide (*χ*² = 18.29, *p* < 0.0001), as well as by all main effects (all *χ*² ³67.07, *p* < 0.001; Supplementary Table S1).

**Figure 1.**
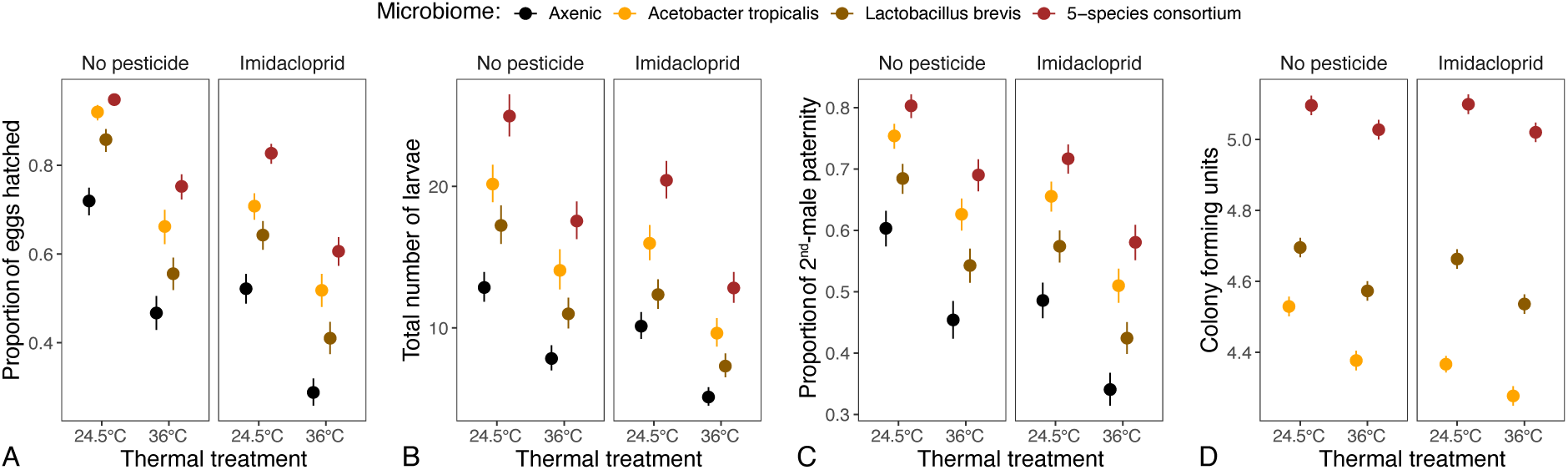
Effects of varying thermal and pesticide conditions and microbiome types on (A) hatching success, (B) male-mediated female fecundity, and (C) competitive fertilization success. (D) Bacterial abundance/colony forming units of different microbial treatments in response to varying thermal and pesticide conditions. Each circle represents the mean of all replicates in that treatment combination with error bars corresponding to 95% confidence intervals.

### Male-mediated fecundity

Next, we investigated male-mediated fecundity, quantified as the total number of larvae produced per female. In contrast to hatching success, the three-way interaction was not significant (*χ*² = 0.68, *df* = 3, *p* = 0.878) for this metric and was removed from the final model. The simplified model revealed significant two-way interactions between microbiome and temperature (*N* = 650, *χ*² = 8.06, df = 3, *p* = 0.042) and between temperature and pesticide (*χ*² = 8.06, df = 1, *p* = 0.005; Fig. 1B, Supplementary Table S2). All main effects were again significant (all *χ*² ³23.92, *p* < 0.0001).

### Sperm competitiveness

Given reports of adverse effects of increased temperatures and pesticide exposure on sperm quality and function (Chirault *et al*. 2015; Meena *et al*. 2024a; Moreira *et al*. 2022; Roy *et al*. 2017; Sales *et al*. 2018, 2024), we evaluated the combined effects of our treatments on the proportion of progeny sired by focal males relative to their standard competitors (i.e., P_2_). Although none of the interactions were statistically significant and thus removed from the model (*N* = 331, all *χ*² = ≤5.26, df = 3, *p* ≥ 0.15), all experimental treatments exhibited highly significant main effects on P_2_ (all *χ*² ³99.60, *p* £0.0001; Fig. 1C, Supplementary Table S3).

### Bacterial abundance in males in response to heat stress and pesticide exposure

We also analyzed the effect of heat stress, pesticide exposure and their interactions on the growth and viability of commensal bacteria inside their host under the different stress treatments by quantifying their colony forming units in homogenates of focal males immediately after mating. For total CFU counts, we found a significant three-way interaction (*N* = 120, *F*_2,106_ = 4.78, *p* = 0.010; Fig. 1D, Supplementary Table S4), suggesting that the bacterial strains differed in their accumulation patterns in adult flies depending on the combined effects of stress treatments. Additionally, we documented significant two-way interactions and main effects of all three factors (all *F* ³10.54, *p* £0.001)

A PERMANOVA controlling for block effects (*F*_1,35_ = 0.20, *p* = 0.874) revealed that temperature was also the dominant driver of variation in the relative abundances of the five bacterial species constituting the consortium, explaining 26.3% of the variation in community composition (*F*_1,35_ = 13.88, *p* = 0.0001; Supplementary Table S5). In contrast, the main effect of pesticide was not statistically significant (2.2% of variation, *F*_1,35_ = 1.16, *p* = 0.314), and the temperature × pesticide interaction showed a marginal effect (4.8% of variation, *F*_1,35_ = 2.55, *p* = 0.068). NMDS based on Bray-Curtis dissimilarities provided a visual representation of community shifts across treatments. The NMDS ordination (Fig. 2A) indicated that the first axis (NMDS1) predominantly tracked the temperature gradient, while the second axis (NMDS2) aligned more, but only weakly, with pesticide exposure. Overlaying species vectors revealed notable differences in species associations (all *R*^2^ ³0.54, *p* £0.0001), in that both *Lactobacillus* species showed positive (³0.68), and all *Acetobacter* exhibited negative (£-0.38), loadings on NMDS1 (Supplementary Table S6), suggesting an increase in the relative abundance of *Acetobacter* in the high-temperature regimes. By contrast, loadings on NMDS2 were negative for *A. tropicalis* (-0.92) and *L. brevis* (-0.44) but positive for all remaining species (³0.56). This axis tended to be biased toward negative values in the absence of imidacloprid, but more ambiguous in its presence.

**Figure 2.**
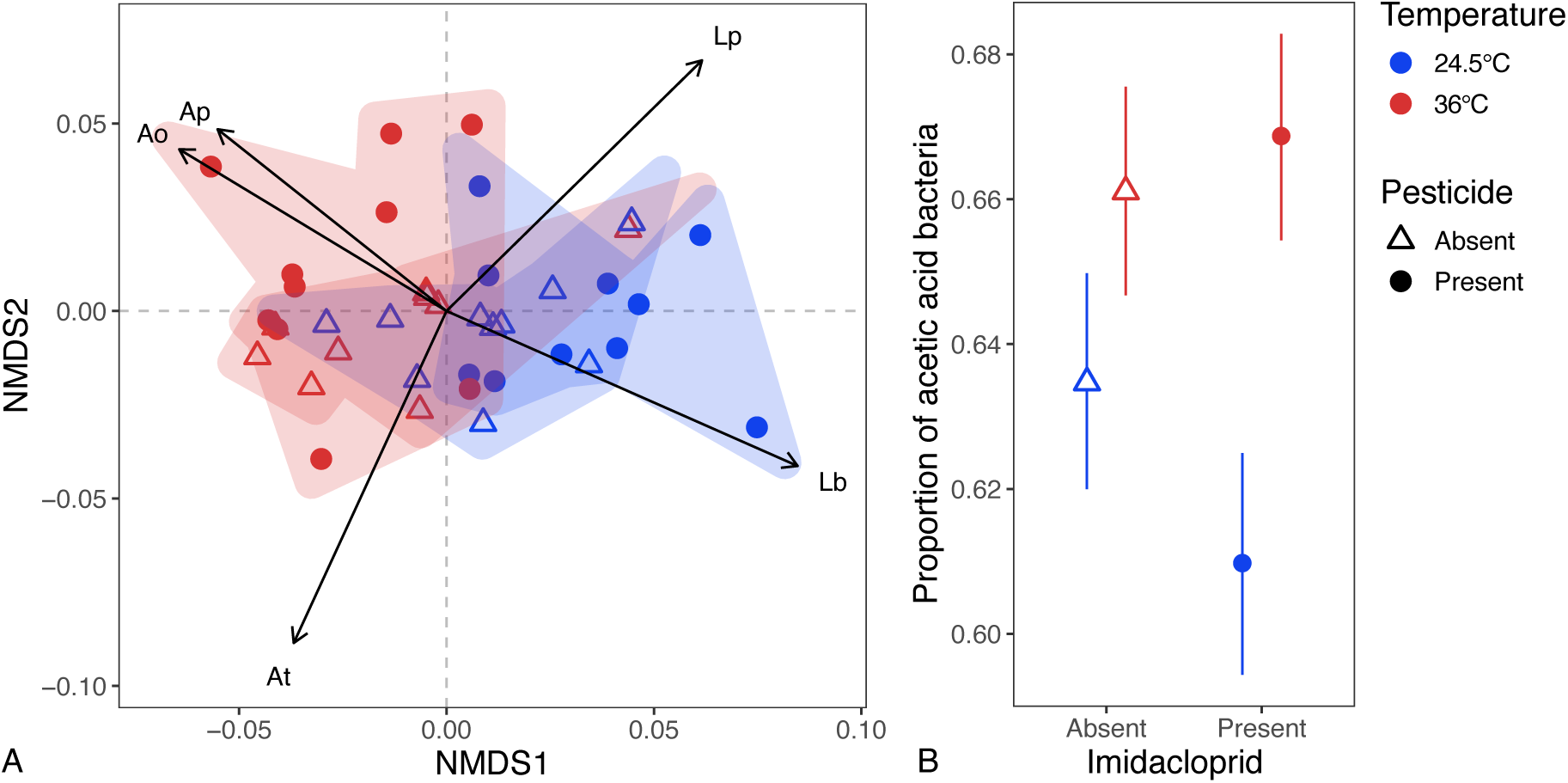
Bacterial abundances in the five-species consortium in response to thermal and pesticide treatments. A: Non-metric multidimensional scaling (NMDS) ordination (Bray-Curtis distance, *k* = 4, stress = 0.046) of microbial community composition across temperature and pesticide treatments. Shaded convex hulls denote 95% confidence areas for each temperature × pesticide group, and arrows the direction and strength of species (Hellinger-transformed) most strongly correlated with the ordination space, as identified by *envfit* (species abbreviations: Ao = *Acetobacter orientalis*, Ap = *A. pasteuriensis*, At = *A. tropicalis*, Lb = *Lactobacillus brevis*, Lp = *L. plantarum*). B: Proportion of acetic acid bacteria in the five-species consortium in response to heat and pesticide stress. Each circle represents the mean of all replicates in that treatment combination with error bars corresponding to 95% confidence intervals.

Finally, in a binomial GLMM focusing only on the relative abundances of acetic acid versus lactic acid bacteria, where the three *Acetobacter* species should theoretically constitute 60% of the total CFUs among the five species, temperature and pesticide presence interacted significantly (*N* = 40, *z* = 2.15, *p* = 0.032). Specifically, the relative representation of *Acetobacter* was increased to around 66% under heat stress (*z* = 2.84, *p* = 0.013), whereas pesticide exposure reduced its representation (*z* = -2.32, *p* = 0.021), although only at the benign temperature (Fig. 3B).

**Figure 3.**
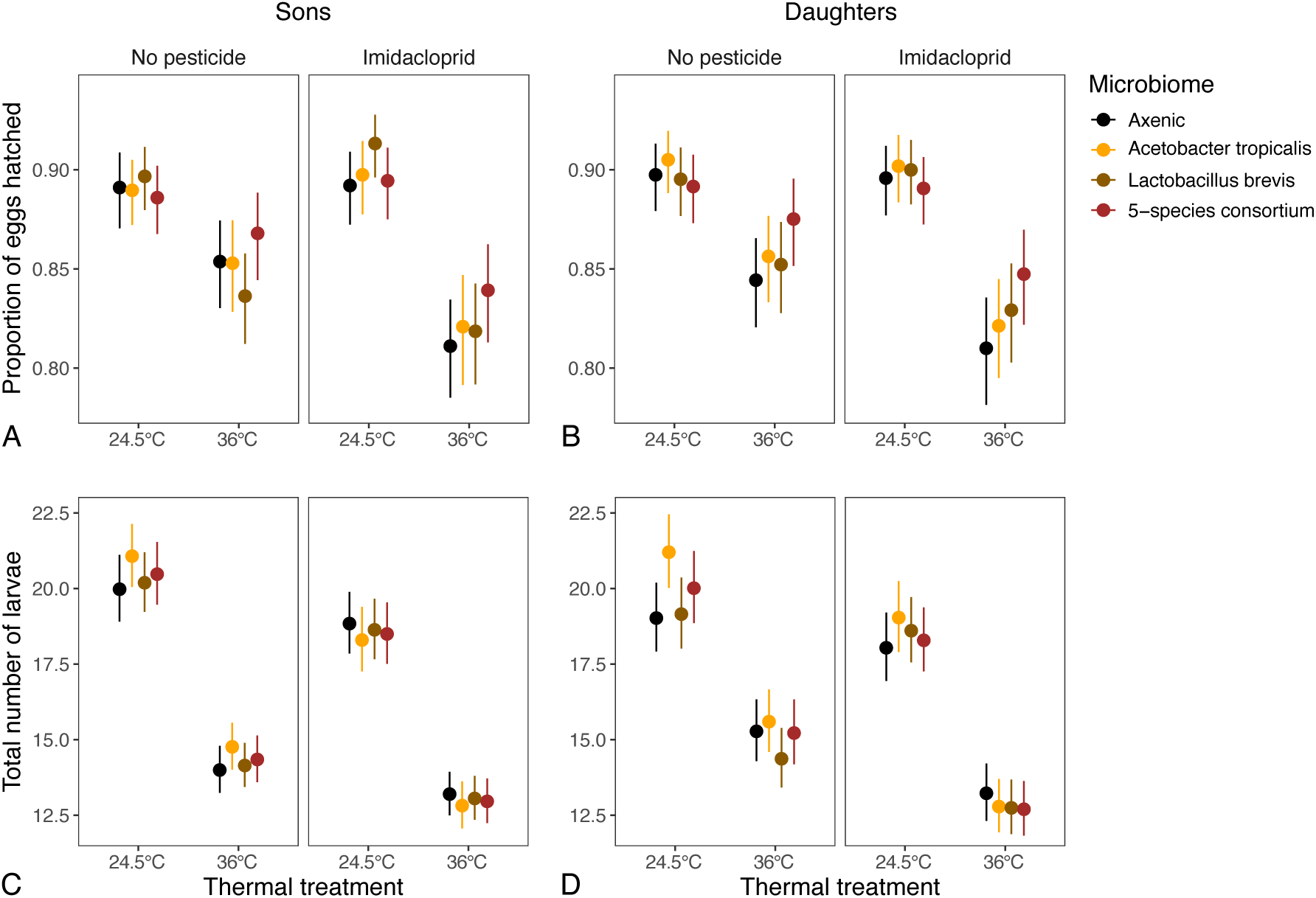
Fitness consequences of paternal thermal and pesticide treatments on fitness measures in male (A, C) and female (B, D) offspring. Panels A and B depict the proportion of eggs hatched out of total number of eggs fertilized, panels C and D the fecundity either of the sons’ mates (C) or of the daughters (D). Each circle represents the mean of all the replicates in that treatment combination with error bars corresponding to 95% confidence intervals.

### Cross-generational effects on offspring fitness

Finally, we investigated how paternal stress exposure affects the next generation by measuring the hatching success and fecundity of F_1_ offspring produced by F_0_ males from each treatment group. We found a significant interactive effect between paternal temperature and pesticide treatments on the hatching success of both male (*N* = 725, *χ*² = 9.45, df = 1, *p* = 0.002; Supplementary Table S7) and female offspring (*N* = 740, *χ*² = 4.73, df = 1, *p* = 0.030; Supplementary Table S8, Fig. 3A,B), indicating that the effects of paternal stress exposure were transmitted to the next generation. However, the paternal microbiome contributed only to very weak and non-statistically significant trends for an interaction with temperature in either offspring sex (*χ*² £6.38, df = 3, *p* = 0.10), suggesting limited cross-generational effects on F_2_ hatching.

The interactive effect of paternal temperature and pesticide exposure was also significant for the fecundity of F_1_ daughters (*N* = 740, *χ*² = 6.11, df = 1, *p* = 0.013), along with a main effect of the paternal microbiome composition (*χ*² = 8.91, df = 3, *p* = 0.030; Fig. 3, Supplementary Table S9). By contrast, the paternal effects on larval numbers in sons was limited to main effects of temperature and pesticide exposure (*χ*² ≥ 26.84, df = 1, *p* < 0.0001), with no contribution of the paternal microbiome (*χ*² = 0.52, df = 3, *p* = 0.915; Fig. 3, Supplementary Table S10).

## Discussion

We investigated the role of the gut microbiome in modulating male reproductive success in *D. melanogaster* under individual and combined exposure to acute heat stress and the pesticide imidacloprid. Our results confirm and extend previous work demonstrating the detrimental effects of sublethal temperatures (Canal Domenech & Fricke 2022; David *et al*. 2005; Fasolo & Krebs 2004; Gandara & Drummond-Barbosa 2022, 2023; Meena *et al*. 2024a; Pilakouta & Baillet 2022; Sales *et al*. 2024; Walsh *et al*. 2019) or imidacloprid (Charpentier *et al*. 2014; Xiao *et al*. 2016; Zhang *et al*. 2021b; Zhou *et al*. 2017) on reproductive functions. Importantly, our study revealed both the combined effects of these stressors and the critical role of the microbiome in mediating the response of stressed males, highlighting the importance of the gut microbiome in mitigating fitness losses. *Acetobacter* generally provided stronger protection than *Lactobacillus*, especially in males with a complex microbial community. Furthermore, we demonstrated that paternal stress exposure imposes fitness costs on the next generation, although these cross-generational effects were not significantly mediated by the paternal microbiome under our experimental conditions designed to minimize direct microbial transmission.

### Interactive effects of combined stressors on male reproduction

Our study revealed that initial exposure to imidacloprid followed by heat shock reduced male reproductive functions to a greater extent than either stressor alone, consistent with previous studies indicating complex, often order-dependent interactions between thermal stress and pesticide toxicity (Ahmed *et al*. 2023; Deng *et al*. 2016; Meng *et al*. 2022; Ricupero *et al*. 2020; Wang *et al*. 2021). While heat stress has been shown to increase the tolerance to subsequent insecticide exposure in diverse insect pests (Cao *et al*. 2019; Ge *et al*. 2013; Gu *et al*. 2010), the reverse order of stress exposure can have opposing effects (Delnat *et al*. 2019; Wang *et al*. 2021), emphasizing the importance of stress sequence in cross-protection.

Pre-exposure to insecticides can also decrease heat stress tolerance, suggesting that warmer temperatures may exacerbate insecticide toxicity depending on the exposure sequence. Increased imidacloprid toxicity at higher temperatures has been documented in mayflies (Macaulay *et al*. 2020), earthworms and collembolans (Bandeira *et al*. 2020), and the terrestrial ectotherm *Hypogastrura viatica* (Silva *et al*. 2023). Our study supports these findings, showing decreased male reproductive success when pesticide exposure preceded heat stress. These multi-stressor studies highlight the complex temperature-toxicity interplay, which can be positive or negative depending on the insecticide mode of action, exposure route, and insect species. Therefore, the sequence of exposure must be carefully considered when evaluating combined stressor effects on insect performance.

The mechanisms underlying these sequence-dependent interactions likely involve intricate physiological adjustments. Exposure to one stressor might compromise pathways needed to cope with the subsequent stressor (lack of cross-tolerance) or trigger shared signaling pathways that amplify susceptibility (negative cross-talk). Resource allocation trade-offs are also probable (Sokolova 2013), where coping with pesticide toxicity may deplete energy reserves or specific molecules (e.g., antioxidants, chaperones) needed for an effective heat stress response. Generally, stressful environments can alter resource allocation among essential processes like growth, reproduction or immune defense to optimize fitness outcomes (Kooĳman 2000). As such, elevated temperature may increase somatic maintenance costs (Verberk *et al*. 2016), potentially reducing resources for defense and repair (Hallman & Brooks 2015).

While the precise mechanisms underlying increased toxicity at high temperatures remain unclear, they may overlap with those explaining the increased toxicity of organophosphate pesticides at higher mean temperatures (Noyes *et al*. 2009; Noyes & Lema 2015). Increased temperatures can enhance the activity of detoxification enzymes like cytochrome P450s (Wang *et al*. 2021), accelerating pesticide breakdown (Ahmad 2007). Increased uptake of chemicals at higher temperatures (Holmstrup *et al*. 2010) has been shown to increase membrane permeability (Noyes & Lema 2015) or inhibit growth (Gardner *et al*. 2011), possibly due to higher basal metabolism (Hooper *et al*. 2013). Stressful conditions can promote the accumulation of reactive oxygen species (ROS) due to reduced respiration and increased water loss, thereby negatively affecting development and reproduction (Dampc *et al*. 2021). Low doses of imidacloprid, for example, can cause ROS formation and metabolic impairment in *D. melanogaster* (Martelli *et al*. 2020). Both pesticide tolerance (Bagchi *et al*. 1996) and heat stress (Neven 2000) are also associated with overexpression of heat-shock proteins. In honeybees, Hsp70 and Hsp90 are upregulated in response to both thermal stress and imidacloprid (Manzi *et al*. 2020). Such chaperone activation could also protect *D. melanogaster* males against combined stressors. If so, the observed fitness loss under both stressors might at least in part be attributable to the costs associated with enhanced Hsp expression (Hoekstra *et al*. 2013; Li *et al*. 2015), which may become overwhelming under combined stress.

### Microbiome as a modulator of stress resilience

A key finding of our study was the significant role of the gut microbiome in mitigating the detrimental reproductive consequences of heat and pesticide stress. Axenic males consistently performed worse than colonized males across all measured fitness components, highlighting the baseline importance of the microbiome for reproduction and its crucial role under stress. The reduced fitness of axenic males under combined stress suggests an impaired ability to metabolize and excrete imidacloprid, potentially leading to higher internal toxin loads and increased vulnerability to subsequent heat stress.

Interestingly, the degree of protection depended on microbial composition. While both *Lactobacillus brevis* and *Acetobacter tropicalis* mono-associations offered benefits over the axenic state, the five-species community consistently provided the greatest resilience, underscoring the potential importance of microbial diversity and inter-species interactions for robust host function (Coyte *et al*. 2015; Freilich *et al*. 2011; Levy & Borenstein 2013; Magnúsdóttir *et al*. 2017; Noecker *et al*. 2019; Zelezniak *et al*. 2015). Mono-associations might lack the metabolic versatility or synergistic cross-feeding capabilities present in more complex consortia, limiting their ability to buffer the host against multiple stressors.

Our results partially contrast with Daisley *et al*. (2017), who reported that *Lactobacillus plantarum* (Lp39) supplementation rescued *D. melanogaster* from harmful imidacloprid effects, accompanied by increased *Acetobacter* and *Lactobacillus* abundance. We did not observe such a strong rescue with *L. brevis*, nor an increased abundance of either bacterial genus. This discrepancy could stem from strain-specific functions, similar to opposing effects on chlorpyrifos (CP) toxicity between *L. plantarum* ISO and *Lactobacillus ATCC* in *D. melanogaster* (Daisley *et al*. 2017), or differences in experimental protocols (e.g., our acute 18h imidacloprid exposure versus Daisley *et al*.’s (2017) continued exposure).

We also observed shifts in the relative abundance of *Acetobacter* and *Lactobacillus* within the host, aligning with studies showing heat-induced microbiome disruptions (Jaramillo & Castañeda 2021; Kokou *et al*. 2018; Moghadam *et al*. 2018). *Lactobacillus* appeared more abundant and less affected by treatments in mono-associations, whereas *Acetobacter* seemed to be overrepresented in the five-species consortium relative to the expected proportion of 60%. Generally, we found a higher proportional representation of *Acetobacter* following acute heat stress, consistent with prior work (Moghadam *et al*. 2018). Beyond Moghadam *et al*.’s (2018) thermal treatments throughout development, however, our findings demonstrate that such compositional shifts can occur within hours.

The mechanisms by which the microbiome enhances stress tolerance likely involve multiple pathways. Beyond direct detoxification (Cheng *et al*. 2017; Jones *et al*. 2015; Kikuchi *et al*. 2012; Pang *et al*. 2018), gut microbes significantly impact host nutrition and metabolism (Douglas 2009; Ridley *et al*. 2012), which are intrinsically linked to thermal tolerance in ectotherms (Henry *et al*. 2020; Moeller *et al*. 2020; Moghadam *et al*. 2018; Semsar-Kazerouni *et al*. 2020). In *D. melanogaster*, the gut microbiota influences nutrient allocation and the thermo-protectant trehalose (Overgaard *et al*. 2007; Ridley *et al*. 2012; Schwasinger-Schmidt

*et al*. 2012). It can also modulate host immune responses and potentially Hsp expression in the gut epithelium (Arnal & Lalles 2016; Liu *et al*. 2014). Given the critical role of Hsps in mitigating thermal stress in *D. melanogaster* (Calabria *et al*. 2012; Sørensen *et al*. 2003), and their rapid induction after heat exposure (Hoekstra *et al*. 2013), the microbiota may enhance heat tolerance by promoting higher Hsp levels. Disrupting the microbiome can alter these processes, as seen with heat-induced reductions in *Buchnera aphidicola* in pea aphids lowering heat resistance (Dunbar *et al*. 2007), and temperature-driven microbiome shifts increasing imidacloprid susceptibility in *Nilaparvata lugens* (Zhang *et al*. 2021a).

### Cross-generational impacts of parental stress

We found clear evidence for cross-generational fitness costs of paternal stress exposure, with reduced hatching success and fecundity in offspring of stressed fathers. Cross-generational effects can either increase or decrease offspring vulnerability to stressors (Brausch & Salice 2011; Kim *et al*. 2014; Pölkki *et al*. 2012; Reátegui-Zirena *et al*. 2017; Schultz *et al*. 2016). Our results corroborate studies reporting detrimental cross- or transgenerational effects of paternal heat stress (Ferrer *et al*. 2013; Guillaume *et al*. 2016; Shama *et al*. 2014; Walsh *et al*. 2014) or toxicant exposure (Bhandari *et al*. 2015; Kimberly & Salice 2015; Schultz *et al*. 2016; Yu & Liao 2016), possibly through epigenetic processes (Schultz *et al*. 2016). The observed synergism between paternal heat and pesticide exposure in offspring fitness underscores the potential for long-lasting consequences of multi-stressor events.

Offspring from stressed fathers exhibited reduced fecundity and hatching success, indicating that paternal exposure does not buffer against these environmental stressors. Our findings support the view that cross-generational effects of stress are often non-adaptive or costly for offspring (Uller *et al*. 2013), highlighting the potential for cumulative fitness declines across generations in populations facing recurrent multi-stressor events.

Intriguingly, the paternal microbiome did not significantly mediate these cross-generational effects in our study. This could be attributable to our experimental design, which deliberately minimized vertical transmission of microbes by using axenic mothers and transferring these and their offspring to sterile food vials immediately after mating or hatching, respectively. This allowed us to isolate direct paternal effects (potentially epigenetic or related to gamete quality) based on the sire’s microbial status. However, we cannot rule out a role for the paternal microbiome if transmission occurs naturally (e.g., during copulation) (Guilhot *et al*. 2025), potentially triggering differential maternal effects or microbial deposition on eggs (direct paternal transmission of microbes to their offspring seems rather unlikely as males have little control over female oviposition). Future studies explicitly allowing or manipulating vertical transmission would be needed to explore this.

### Conclusions and broader implications

In conclusion, our study demonstrates that acute heat stress and sublethal pesticide exposure interact, often synergistically, to negatively impact male reproductive fitness in *D. melanogaster*, with consequences extending to offspring viability and fecundity. Crucially, we show that the gut microbiome is a key determinant of resilience, offering significant protection against these stressors, with benefits enhanced by microbial complexity. However, the microbiome itself is vulnerable to disruption by these same stressors.

Our findings underscore the critical need to consider multi-stressor interactions and the role of host-associated microbes when assessing the ecological risks posed by climate change and agrochemical use. The observed negative cross-generational effects further highlight the potential vulnerability of insect populations, particularly those with depleted or altered microbiomes. Future research should explore these interactions across diverse species, environments, and stressor combinations to improve ecological forecasting and develop strategies for mitigating biodiversity loss in the Anthropocene.

## Acknowledgements

We thank W. Ludington for providing the bacterial strains and T. Ratz and G. Raina for valuable comments on our manuscript. This project was funded by the Swiss National Science Foundation (grant PP00P3_202662 to S.L.) and a University of Zurich Postdoctoral Fellowship (FK-23-127 to K.M.).

## Author contributions

KM and SL conceived the study and acquired funding; KM, AM, ANDN, BE, SHS and RATS collected the data; KM and SL conducted statistical analyses and drafted the manuscript, which was reviewed and approved by all other authors.

## Supplementary material

**Table S1:** Results from a generalized linear model on hatching success. The model assessed the impact of male microbiome (axenic, *Acetobacter tropicalis*, *Lactobacillus brevis*, control), temperature (24.5°C/36°C), and pesticide exposure (absent/present). Total sample size: *N =* 650.

| *Predictor* | *df* | $\chi^2$ | *P* |
| --- | --- | --- | --- |
| Microbiome | 3 | 215.696 | **<0.001** |
| Temperature | 1 | 93.332 | **<0.001** |
| Pesticide | 1 | 67.065 | **<0.001** |
| Block | 1 | 6.955 | **0.008** |
| Microbiome × Temperature | 3 | 22.007 | **<0.001** |
| Microbiome × Pesticide | 3 | 18.286 | **<0.001** |
| Temperature × Pesticide | 1 | 0.292 | 0.589 |
| Microbiome × Temperature × Pesticide | 3 | 15.190 | **0.002** |

Coefficients of the model described above.
| *Term* | *Estimate* | *SE* | *z* | *P* |
| --- | --- | --- | --- | --- |
| (Intercept) | 0.889 | 0.082 | 10.847 | **<0.001** |
| Microbiome[At] | 1.501 | 0.141 | 10.624 | **<0.001** |
| Microbiome[Lb] | 0.855 | 0.133 | 6.442 | **<0.001** |
| Microbiome[Control] | 1.960 | 0.154 | 12.713 | **<0.001** |
| Temperature[36°C] | -1.074 | 0.111 | -9.661 | **<0.001** |
| Pesticide[Present] | -0.855 | 0.104 | -8.189 | **<0.001** |
| Block[2] | 0.107 | 0.040 | 2.637 | **0.008** |
| Microbiome[At] × Temperature[36°C] | -0.698 | 0.184 | -3.793 | **<0.001** |
| Microbiome[Lb] × Temperature[36°C] | -0.500 | 0.171 | -2.915 | **0.004** |
| Microbiome[Control] × Temperature[36°C] | -0.717 | 0.189 | -3.796 | **<0.001** |
| Microbiome[At] × Pesticide[Present] | -0.704 | 0.173 | -4.062 | **<0.001** |
| Microbiome[Lb] × Pesticide[Present] | -0.355 | 0.166 | -2.144 | **0.032** |
| Microbiome[Control] × Pesticide[Present] | -0.483 | 0.186 | -2.589 | **0.010** |
| Temperature[36°C] × Pesticide[Present] | 0.082 | 0.151 | 0.540 | 0.589 |
| Microbiome[At] × Temperature[36°C] × Pesticide[Present] | 0.877 | 0.236 | 3.721 | **<0.001** |
| Microbiome[Lb] × Temperature[36°C] × Pesticide[Present] | 0.542 | 0.226 | 2.400 | **0.016** |
| Microbiome[Control] × Temperature[36°C] × Pesticide[Present] | 0.576 | 0.239 | 2.405 | **0.016** |

**Table S2:** Results from a generalized linear model on the total number of larvae produced. The model assessed the impact of male microbiome (axenic, *Acetobacter tropicalis*, *Lactobacillus brevis*, control), temperature (24.5°C/36°C), and pesticide exposure (absent/present). The three-way interaction was removed due to non-significance (*P* = 0.88). Total sample size: *N =* 650.

| *Predictor* | *df* | $\chi^2$ | *P* |
| --- | --- | --- | --- |
| Microbiome | 3 | 207.693 | **<0.001** |
| Temperature | 1 | 85.063 | **<0.001** |
| Pesticide | 1 | 23.921 | **<0.001** |
| Block | 1 | 0.718 | 0.397 |
| Microbiome × Temperature | 3 | 8.223 | **0.042** |
| Microbiome × Pesticide | 3 | 4.264 | 0.234 |
| Temperature × Pesticide | 1 | 8.061 | **0.005** |

Coefficients of the model described above.
| *Term* | *Estimate* | *SE* | *z* | *P* |
| --- | --- | --- | --- | --- |
| (Intercept) | 2.554 | 0.040 | 63.187 | **<0.001** |
| Microbiome[At] | 0.443 | 0.049 | 9.039 | **<0.001** |
| Microbiome[Lb] | 0.275 | 0.053 | 5.206 | **<0.001** |
| Microbiome[Control] | 0.652 | 0.047 | 13.873 | **<0.001** |
| Temperature[36°C] | -0.521 | 0.057 | -9.223 | **<0.001** |
| Pesticide[Present] | -0.260 | 0.053 | -4.891 | **<0.001** |
| Block[2] | 0.018 | 0.021 | 0.848 | 0.397 |
| Microbiome[At] × Temperature[36°C] | 0.152 | 0.069 | 2.190 | **0.029** |
| Microbiome[Lb] × Temperature[36°C] | 0.094 | 0.072 | 1.311 | 0.190 |
| Microbiome[Control] × Temperature[36°C] | 0.175 | 0.064 | 2.724 | **0.006** |
| Microbiome[At] × Pesticide[Present] | 0.021 | 0.066 | 0.318 | 0.750 |
| Microbiome[Lb] × Pesticide[Present] | -0.054 | 0.069 | -0.788 | 0.431 |
| Microbiome[Control] × Pesticide[Present] | 0.065 | 0.062 | 1.042 | 0.297 |
| Temperature[36°C] × Pesticide[Present] | -0.126 | 0.045 | -2.839 | **0.005** |

**Table S3:** Results from a generalized linear model on the proportion of second-male offspring (sperm offense ability). The model assessed the impact of male microbiome (axenic, *Acetobacter tropicalis*, *Lactobacillus brevis*, control), temperature (24.5°C/36°C), and pesticide exposure (absent/present). All interactions were sequentially removed due to non-significance (*P* > 0.15). Total sample size: *N =* 331.

| *Predictor* | *df* | $\chi^2$ | *P* |
| --- | --- | --- | --- |
| Microbiome | 3 | 218.453 | **<0.001** |
| Temperature | 1 | 162.250 | **<0.001** |
| Pesticide | 1 | 99.596 | **<0.001** |
| Block | 1 | 0.827 | 0.363 |

Coefficients of the model described above.
| *Term* | *Estimate* | *SE* | *z* | *P* |
| --- | --- | --- | --- | --- |
| (Intercept) | 0.397 | 0.067 | 5.902 | **<0.001** |
| Microbiome[At] | 0.700 | 0.068 | 10.320 | **<0.001** |
| Microbiome[Lb] | 0.356 | 0.068 | 5.252 | **<0.001** |
| Microbiome[Control] | 0.985 | 0.072 | 13.716 | **<0.001** |
| Temperature[36°C] | -0.604 | 0.047 | -12.738 | **<0.001** |
| Pesticide[Present] | -0.476 | 0.048 | -9.980 | **<0.001** |
| Block[2] | 0.043 | 0.048 | 0.910 | 0.363 |

**Table S4:** Results from a linear model on colony forming units. The model assessed the impact of male microbiome (axenic, *Acetobacter tropicalis*, *Lactobacillus brevis*, control), temperature (24.5°C/36°C), and pesticide exposure (absent/present). Total sample size: *N =* 120.

| *Predictor* | *df* | *F* | *P* |
| --- | --- | --- | --- |
| Temperature | 1 | 135.611 | **<0.001** |
| Pesticide | 1 | 146.557 | **<0.001** |
| Microbiome | 2 | 991.897 | **<0.001** |
| Block | 2 | 0.156 | 0.856 |
| Temperature × Pesticide | 1 | 11.281 | **0.001** |
| Temperature × Microbiome | 2 | 10.543 | **<0.001** |
| Pesticide × Microbiome | 2 | 43.073 | **<0.001** |
| Temperature × Pesticide × Microbiome | 2 | 4.778 | **0.010** |
| Residuals | 106 |  |  |

Coefficients of the model described above.
| *Term* | *Estimate* | *SE* | *t* | *P* |
| --- | --- | --- | --- | --- |
| (Intercept) | 4.530 | 0.010 | 470.489 | **<0.001** |
| Temperature[36°C] | -0.152 | 0.013 | -11.645 | **<0.001** |
| Pesticide[Present] | -0.163 | 0.013 | -12.106 | **<0.001** |
| Microbiome[Lb] | 0.166 | 0.013 | 12.706 | **<0.001** |
| Microbiome[Control] | 0.567 | 0.013 | 43.323 | **<0.001** |
| Block[2] | 0.003 | 0.005 | 0.510 | 0.611 |
| Block[3] | -0.006 | 0.031 | -0.187 | 0.852 |
| Temperature[36°C] × Pesticide[Present] | 0.063 | 0.019 | 3.359 | **0.001** |
| Temperature[36°C] × Microbiome[Lb] | 0.030 | 0.018 | 1.624 | 0.107 |
| Temperature[36°C] × Microbiome[Control] | 0.084 | 0.018 | 4.532 | **<0.001** |
| Pesticide[Present] × Microbiome[Lb] | 0.130 | 0.019 | 6.944 | **<0.001** |
| Pesticide[Present] × Microbiome[Control] | 0.166 | 0.019 | 8.851 | **<0.001** |
| Temperature[36°C] × Pesticide[Present] × Microbiome[Lb] | -0.067 | 0.026 | -2.559 | **0.012** |
| Temperature[36°C] × Pesticide[Present] × Microbiome[Control] | -0.074 | 0.026 | -2.792 | **0.006** |

**Table S5:**
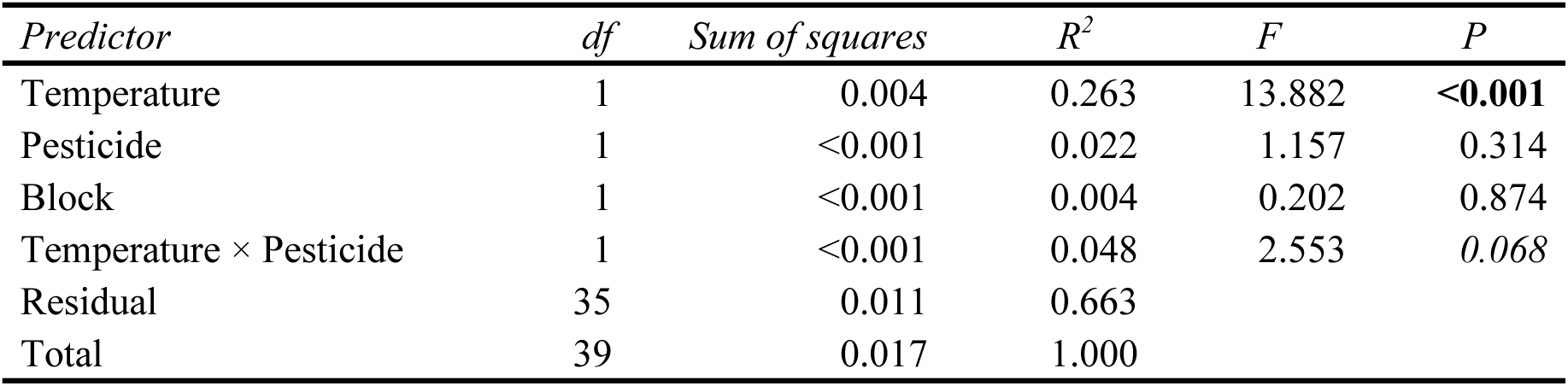
Results of a PERMANOVA examining the temperature and pesticide effects on relative bacterial abundances (based on colony-forming units) in the five-species consortium.

**Table S6:** Ordination of the Bray-Curtis dissimilarities between the relative abundances of the five bacterial species via non-metric multidimensional scaling (NMDS).

| *Species* | *NMDS1* | *NMDS2* | *R*<sup>2</sup> | *P* |
| --- | --- | --- | --- | --- |
| *A. tropicalis* | -0.384 | -0.923 | 0.920 | **<0.001** |
| *A. orientalis* | -0.831 | 0.556 | 0.601 | **<0.001** |
| *A. pasteurianus* | -0.751 | 0.660 | 0.539 | **<0.001** |
| *L. brevis* | 0.899 | -0.439 | 0.887 | **<0.001** |
| *L. plantarum* | 0.678 | 0.735 | 0.826 | **<0.001** |

**Table S7:** Results from a binomial generalized linear mixed-effects model on hatching success among the eggs laid by the mates of the sons. The model assessed the impact of male microbiome (axenic, *Acetobacter tropicalis*, *Lactobacillus brevis*, control), temperature (24.5°C/36°C), and pesticide exposure (absent/present), incorporating sire to account for non-independence among siblings. The three-way interaction was removed due to non-significance (*P* = 0.48). Total sample size: *N =* 725.

| *Predictor* | *df* | $\chi^2$ | *P* |
| --- | --- | --- | --- |
| Microbiome | 3 | 0.903 | 0.825 |
| Temperature | 1 | 8.359 | **0.004** |
| Pesticide | 1 | 0.008 | 0.929 |
| Block | 1 | 5.924 | **0.015** |
| Microbiome × Temperature | 3 | 6.380 | 0.095 |
| Microbiome × Pesticide | 3 | 1.681 | 0.641 |
| Temperature × Pesticide | 1 | 9.451 | **0.002** |

Random effects variance components from the model described above.
| *Group* | *Term* | *SD* | *Variance* |
| --- | --- | --- | --- |
| Sire | (Intercept) | 0 | 0 |

Fixed effects estimates from the model described above.
| *Term* | *Estimate* | *SE* | *z* | *P* |
| --- | --- | --- | --- | --- |
| (Intercept) | 2.163 | 0.103 | 21.076 | **<0.001** |
| Microbiome[At] | -0.014 | 0.126 | -0.113 | 0.910 |
| Microbiome[Lb] | 0.059 | 0.127 | 0.467 | 0.640 |
| Microbiome[Control] | -0.052 | 0.127 | -0.406 | 0.685 |
| Temperature[36°C] | -0.337 | 0.117 | -2.891 | **0.004** |
| Pesticide[Present] | 0.010 | 0.118 | 0.089 | 0.929 |
| Block[2] | -0.125 | 0.051 | -2.434 | **0.015** |
| Microbiome[At] × Temperature[36°C] | 0.008 | 0.147 | 0.054 | 0.957 |
| Microbiome[Lb] × Temperature[36°C] | -0.192 | 0.143 | -1.340 | 0.180 |
| Microbiome[Control] × Temperature[36°C] | 0.170 | 0.146 | 1.165 | 0.244 |
| Microbiome[At] × Pesticide[Present] | 0.072 | 0.147 | 0.489 | 0.625 |
| Microbiome[Lb] × Pesticide[Present] | 0.183 | 0.142 | 1.283 | 0.199 |
| Microbiome[Control] × Pesticide[Present] | 0.076 | 0.146 | 0.524 | 0.600 |
| Temperature[36°C] × Pesticide[Present] | -0.317 | 0.103 | -3.074 | **0.002** |

**Table S8:** Results from a binomial generalized linear mixed-effects model on hatching success among the eggs laid by the daughters. The model assessed the impact of male microbiome (axenic, *Acetobacter tropicalis*, *Lactobacillus brevis*, control), temperature (24.5°C/36°C), and pesticide exposure (absent/present), incorporating sire to account for non-independence among siblings. The three-way interaction was removed due to non-significance (*P* = 0.87). Total sample size: *N =* 740.

| *Predictor* | *df* | $\chi^2$ | *P* |
| --- | --- | --- | --- |
| Microbiome | 3 | 1.500 | 0.682 |
| Temperature | 1 | 17.866 | **<0.001** |
| Pesticide | 1 | 0.023 | 0.878 |
| Block | 1 | 7.816 | **0.005** |
| Microbiome × Temperature | 3 | 6.151 | 0.104 |
| Microbiome × Pesticide | 3 | 0.420 | 0.936 |
| Temperature × Pesticide | 1 | 4.733 | **0.030** |

Random effects variance components from the model described above.
| *Group* | *Term* | *SD* | *Variance* |
| --- | --- | --- | --- |
| Sire | (Intercept) | 0 | 0 |

Fixed effects estimates from the model described above.
| *Term* | *Estimate* | *SE* | *z* | *P* |
| --- | --- | --- | --- | --- |
| (Intercept) | 2.240 | 0.097 | 23.120 | **<0.001** |
| Microbiome[At] | 0.085 | 0.126 | 0.675 | 0.499 |
| Microbiome[Lb] | -0.024 | 0.127 | -0.188 | 0.851 |
| Microbiome[Control] | -0.062 | 0.126 | -0.496 | 0.620 |
| Temperature[36°C] | -0.478 | 0.113 | -4.227 | **<0.001** |
| Pesticide[Present] | -0.018 | 0.116 | -0.153 | 0.878 |
| Block[2] | -0.143 | 0.051 | -2.796 | **0.005** |
| Microbiome[At] × Temperature[36°C] | 0.009 | 0.144 | 0.063 | 0.950 |
| Microbiome[Lb] × Temperature[36°C] | 0.085 | 0.144 | 0.591 | 0.555 |
| Microbiome[Control] × Temperature[36°C] | 0.319 | 0.146 | 2.184 | **0.029** |
| Microbiome[At] × Pesticide[Present] | -0.019 | 0.143 | -0.133 | 0.894 |
| Microbiome[Lb] × Pesticide[Present] | 0.069 | 0.144 | 0.480 | 0.631 |
| Microbiome[Control] × Pesticide[Present] | 0.008 | 0.146 | 0.054 | 0.957 |
| Temperature[36°C] × Pesticide[Present] | -0.223 | 0.103 | -2.176 | **0.030** |

**Table S9:**
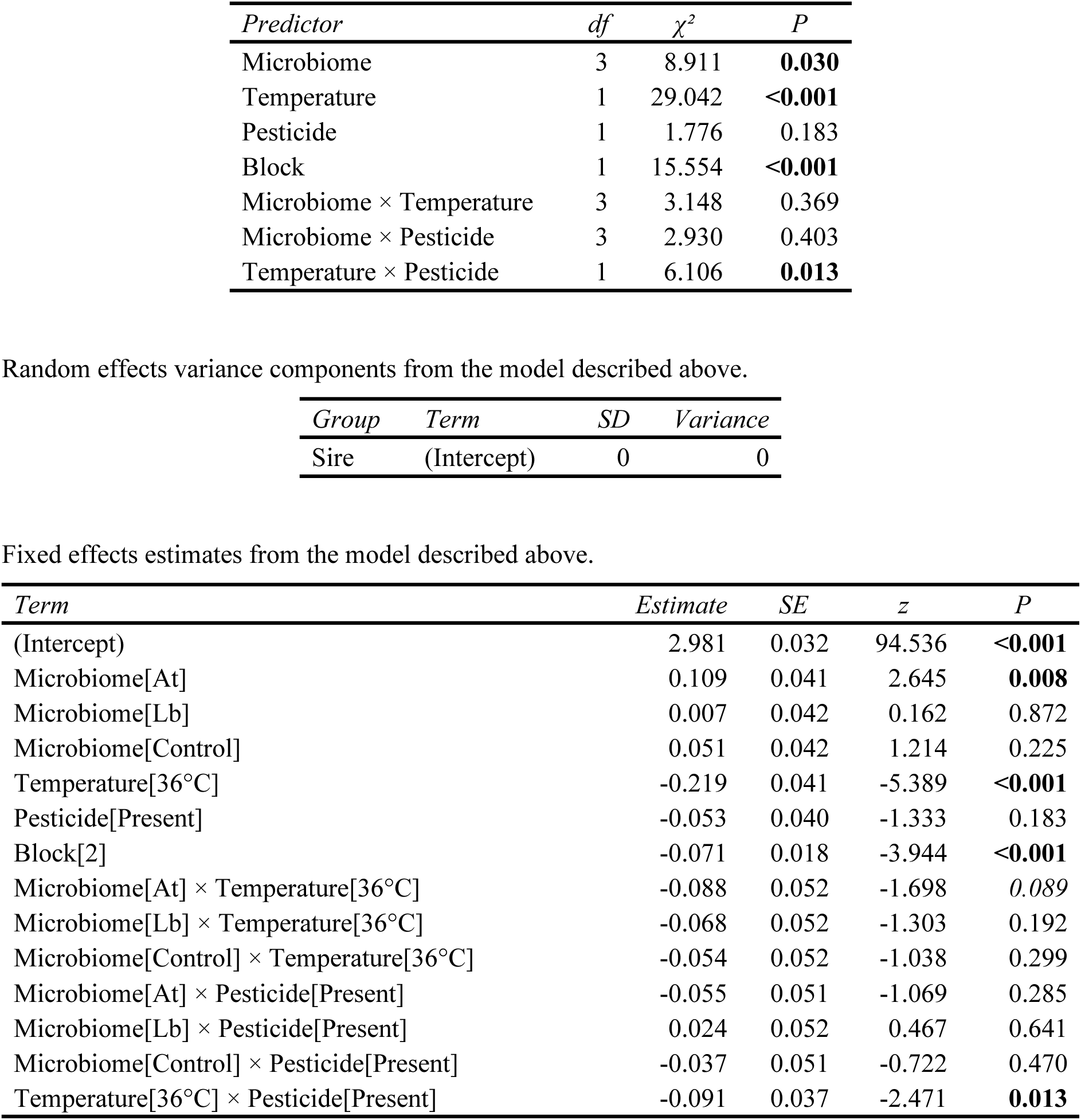
Results from a Poisson generalized linear mixed-effects model on the total number of larvae produced by the daughters. The model assessed the impact of male microbiome (axenic, *Acetobacter tropicalis*, *Lactobacillus brevis*, control), temperature (24.5°C/36°C), and pesticide exposure (absent/present), incorporating sire to account for non-independence among siblings. The three-way interaction was removed due to non-significance (*P* = 0.22). Total sample size: *N =* 740.

| *Predictor* | *df* | $\chi^2$ | *P* |
| --- | --- | --- | --- |
| Microbiome | 3 | 8.911 | **0.030** |
| Temperature | 1 | 29.042 | **<0.001** |
| Pesticide | 1 | 1.776 | 0.183 |
| Block | 1 | 15.554 | **<0.001** |
| Microbiome × Temperature | 3 | 3.148 | 0.369 |
| Microbiome × Pesticide | 3 | 2.930 | 0.403 |
| Temperature × Pesticide | 1 | 6.106 | **0.013** |

Random effects variance components from the model described above.
| *Group* | *Term* | *SD* | *Variance* |
| --- | --- | --- | --- |
| Sire | (Intercept) | 0 | 0 |

Fixed effects estimates from the model described above.
| *Term* | *Estimate* | *SE* | *z* | *P* |
| --- | --- | --- | --- | --- |
| (Intercept) | 2.981 | 0.032 | 94.536 | **<0.001** |
| Microbiome[At] | 0.109 | 0.041 | 2.645 | **0.008** |
| Microbiome[Lb] | 0.007 | 0.042 | 0.162 | 0.872 |
| Microbiome[Control] | 0.051 | 0.042 | 1.214 | 0.225 |
| Temperature[36°C] | -0.219 | 0.041 | -5.389 | **<0.001** |
| Pesticide[Present] | -0.053 | 0.040 | -1.333 | 0.183 |
| Block[2] | -0.071 | 0.018 | -3.944 | **<0.001** |
| Microbiome[At] × Temperature[36°C] | -0.088 | 0.052 | -1.698 | 0.089 |
| Microbiome[Lb] × Temperature[36°C] | -0.068 | 0.052 | -1.303 | 0.192 |
| Microbiome[Control] × Temperature[36°C] | -0.054 | 0.052 | -1.038 | 0.299 |
| Microbiome[At] × Pesticide[Present] | -0.055 | 0.051 | -1.069 | 0.285 |
| Microbiome[Lb] × Pesticide[Present] | 0.024 | 0.052 | 0.467 | 0.641 |
| Microbiome[Control] × Pesticide[Present] | -0.037 | 0.051 | -0.722 | 0.470 |
| Temperature[36°C] × Pesticide[Present] | -0.091 | 0.037 | -2.471 | **0.013** |

**Table S10:**
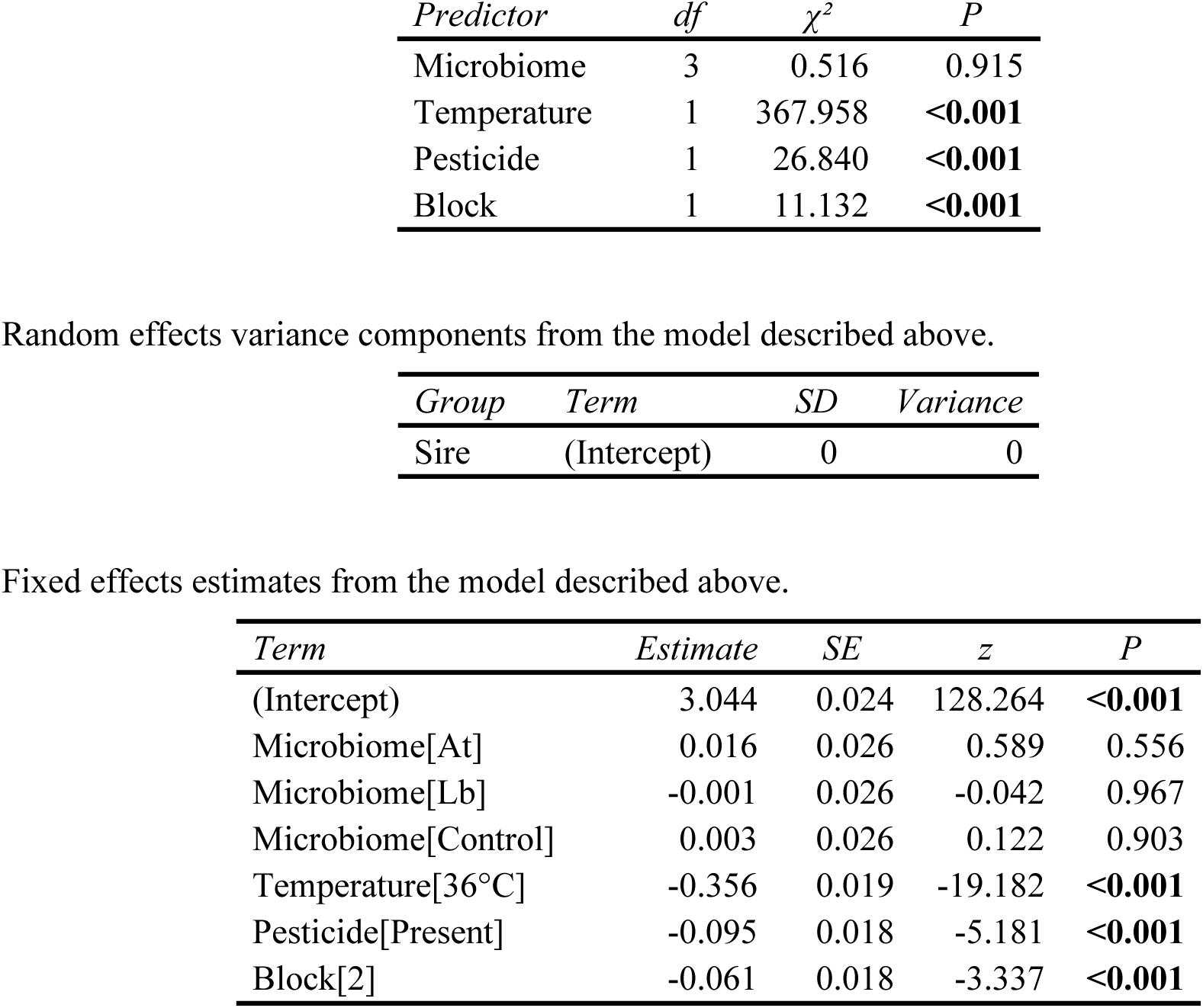
Results from a Poisson generalized linear mixed-effects model on the total number of larvae produced by the mates of the sons. The model assessed the impact of male microbiome (axenic, *Acetobacter tropicalis*, *Lactobacillus brevis*, control), temperature (24.5°C/36°C), and pesticide exposure (absent/present), incorporating sire to account for non-independence among siblings. The interactions were sequentially removed due to non-significance (*P* ≥ 0.20). Total sample size: *N =* 725.

| *Predictor* | *df* | $\chi^2$ | *P* |
| --- | --- | --- | --- |
| Microbiome | 3 | 0.516 | 0.915 |
| Temperature | 1 | 367.958 | **<0.001** |
| Pesticide | 1 | 26.840 | **<0.001** |
| Block | 1 | 11.132 | **<0.001** |

Random effects variance components from the model described above.
| *Group* | *Term* | *SD* | *Variance* |
| --- | --- | --- | --- |
| Sire | (Intercept) | 0 | 0 |

Fixed effects estimates from the model described above.
| *Term* | *Estimate* | *SE* | *z* | *P* |
| --- | --- | --- | --- | --- |
| (Intercept) | 3.044 | 0.024 | 128.264 | **<0.001** |
| Microbiome[At] | 0.016 | 0.026 | 0.589 | 0.556 |
| Microbiome[Lb] | -0.001 | 0.026 | -0.042 | 0.967 |
| Microbiome[Control] | 0.003 | 0.026 | 0.122 | 0.903 |
| Temperature[36°C] | -0.356 | 0.019 | -19.182 | **<0.001** |
| Pesticide[Present] | -0.095 | 0.018 | -5.181 | **<0.001** |
| Block[2] | -0.061 | 0.018 | -3.337 | **<0.001** |

## References

Ahmad, M. (2007). Insecticide resistance mechanisms and their management in *Helicoverpa armigera* (Hübner)–A review. J Agric Res, 45, 319–335.

Ahmed, T., Masood, H.A., Noman, M., Al-Huqail, A.A., Alghanem, S.M., Khan, M.M., et al. (2023). Biogenic silicon nanoparticles mitigate cadmium (Cd) toxicity in rapeseed (*Brassica napus* L.) by modulating the cellular oxidative stress metabolism and reducing Cd translocation. J. Hazard. Mater., 459, 132070.

Arambourou, H. & Stoks, R. (2015). Combined effects of larval exposure to a heat wave and chlorpyrifos in northern and southern populations of the damselfly *Ischnura elegans*. Chemosphere, 128, 148–154.

Arnal, M.-E. & Lalles, J.-P. (2016). Gut epithelial inducible heat-shock proteins and their modulation by diet and the microbiota. Nutr. Rev., 74, 181–197.

Aulsebrook, L.C., Bertram, M.G., Martin, J.M., Aulsebrook, A.E., Brodin, T., Evans, J.P., et al. (2020). Reproduction in a polluted world: implications for wildlife. Reproduction, 160, R13–R23.

Bagchi, D., Bhaitacharya, G. & Stohs, S. (1996). Production of reactive oxygen species by gastric cells in association with *Helicobacter pylori*. Free Radic. Res., 24, 439–450.

Bandeira, F.O., Alves, P.R.L., Hennig, T.B., Toniolo, T., Natal-da-Luz, T. & Baretta, D. (2020). Effect of temperature on the toxicity of imidacloprid to *Eisenia andrei* and *Folsomia candida* in tropical soils. Environ. Pollut., 267, 115565.

Beketov, M.A., Kefford, B.J., Schäfer, R.B. & Liess, M. (2013). Pesticides reduce regional biodiversity of stream invertebrates. Proc. Natl. Acad. Sci., 110, 11039–11043.

Bhandari, R.K., Vom Saal, F.S. & Tillitt, D.E. (2015). Transgenerational effects from early developmental exposures to bisphenol A or 17α-ethinylestradiol in medaka, *Oryzias latipes*. Sci. Rep., 5, 9303.

Bonduriansky, R., Crean, A.J. & Day, T. (2012). The implications of nongenetic inheritance for evolution in changing environments. Evol. Appl., 5, 192–201.

Brausch, J.M. & Salice, C.J. (2011). Effects of an environmentally realistic pesticide mixture on *Daphnia magna* exposed for two generations. Arch. Environ. Contam. Toxicol., 61, 272–279.

Broderick, N.A. & Lemaitre, B. (2012). Gut-associated microbes of *Drosophila melanogaster*. Gut Microbes, 3, 307–321.

Brooks, M.E., Kristensen, K., Van Benthem, K.J., Magnusson, A., Berg, C.W., Nielsen, A., et al. (2017). glmmTMB balances speed and flexibility among packages for zero-inflated generalized linear mixed modeling.

Calabria, G., Dolgova, O., Rego, C., Castañeda, L., Rezende, E., Balanyà, J., et al. (2012). Hsp70 protein levels and thermotolerance in *Drosophila subobscura*: a reassessment of the thermal co-adaptation hypothesis. J. Evol. Biol., 25, 691–700.

Canal Domenech, B. & Fricke, C. (2022). Recovery from heat-induced infertility—A study of reproductive tissue responses and fitness consequences in male *Drosophila melanogaster*. Ecol. Evol., 12, e9563.

Cao, Y., Yang, H., Li, J., Wang, C., Li, C. & Gao, Y. (2019). Sublethal effects of imidacloprid on the population development of western flower thrips *Frankliniella occidentalis* (Thysanoptera: Thripidae). Insects, 10, 3.

Charpentier, G., Louat, F., Bonmatin, J.-M., Marchand, P.A., Vanier, F., Locker, D., et al. (2014). Lethal and sublethal effects of imidacloprid, after chronic exposure, on the insect model *Drosophila melanogaster*. Environ. Sci. Technol., 48, 4096–4102.

Cheng, D., Guo, Z., Riegler, M., Xi, Z., Liang, G. & Xu, Y. (2017). Gut symbiont enhances insecticide resistance in a significant pest, the oriental fruit fly *Bactrocera dorsalis* (Hendel). Microbiome, 5, 1–12.

Chirault, M., Lucas, C., Goubault, M., Chevrier, C., Bressac, C. & Lécureuil, C. (2015). A combined approach to heat stress effect on male fertility in *Nasonia vitripennis*: from the physiological consequences on spermatogenesis to the reproductive adjustment of females mated with stressed males. PLoS One, 10, e0120656.

Coyte, K.Z., Schluter, J. & Foster, K.R. (2015). The ecology of the microbiome: networks, competition, and stability. Science, 350, 663–666.

Crean, A.J. & Immler, S. (2021). Evolutionary consequences of environmental effects on gamete performance. Philos. Trans. R. Soc. B, 376, 20200122.

Dai, W., Holmstrup, M., Slotsbo, S., Bakker, R., Damgaard, C. & van Gestel, C.A. (2023). Heat stress delays detoxification of phenanthrene in the springtail *Folsomia candida*. Chemosphere, 311, 137119.

Daisley, B.A., Trinder, M., McDowell, T.W., Welle, H., Dube, J.S., Ali, S.N., et al. (2017). Neonicotinoid-induced pathogen susceptibility is mitigated by *Lactobacillus plantarum* immune stimulation in a *Drosophila melanogaster* model. Sci. Rep., 7, 2703.

Dampc, J., Mołoń, M., Durak, T. & Durak, R. (2021). Changes in aphid—plant interactions under increased temperature. Biology, 10, 480.

David, J., Araripe, L., Chakir, M., Legout, H., Lemos, B., Pétavy, G., et al. (2005). Male sterility at extreme temperatures: a significant but neglected phenomenon for understanding *Drosophila* climatic adaptations. J. Evol. Biol., 18, 838–846.

De França, S.M., Breda, M.O., Barbosa, D.R., Araujo, A., Guedes, C.A. & Shields, V. (2017). The sublethal effects of insecticides in insects. Biological Control of Pest and Vector Insects.,10,23–39.

DeCourten, B.M. & Brander, S.M. (2017). Combined effects of increased temperature and endocrine disrupting pollutants on sex determination, survival, and development across generations. Sci. Rep., 7, 9310.

Delnat, V., Janssens, L. & Stoks, R. (2019). Whether warming magnifies the toxicity of a pesticide is strongly dependent on the concentration and the null model. Aquat. Toxicol., 211, 38–45.

Deng, Z., Zhang, F., Wu, Z., Yu, Z. & Wu, G. (2016). Chlorpyrifos-induced hormesis in insecticide-resistant and-susceptible *Plutella xylostella* under normal and high temperatures. Bull. Entomol. Res., 106, 378–386.

Desneux, N., Decourtye, A. & Delpuech, J.-M. (2007). The sublethal effects of pesticides on beneficial arthropods. Annu Rev Entomol, 52, 81–106.

Deutsch, C.A., Tewksbury, J.J., Huey, R.B., Sheldon, K.S., Ghalambor, C.K., Haak, D.C., et al. (2008). Impacts of climate warming on terrestrial ectotherms across latitude. Proc. Natl. Acad. Sci. U. S. A., 105, 6668–6672.

Deutsch, C.A., Tewksbury, J.J., Tigchelaar, M., Battisti, D.S., Merrill, S.C., Huey, R.B., et al. (2018). Increase in crop losses to insect pests in a warming climate. Science, 361, 916– 919.

Dougherty, L.R., Frost, F., Maenpaa, M.I., Rowe, M., Cole, B.J., Vasudeva, R., et al. (2024). A systematic map of studies testing the relationship between temperature and animal reproduction. Ecol. Solut. Evid., 5, e12303.

Douglas, A.E. (2009). The microbial dimension in insect nutritional ecology. Funct. Ecol., 23, 38–47.

Dunbar, H.E., Wilson, A.C.C., Ferguson, N.R. & Moran, N.A. (2007). Aphid thermal tolerance is governed by a point mutation in bacterial symbionts. PLoS Biol., 5, e96.

Fasolo, A.G. & Krebs, R.A. (2004). A comparison of behavioural change in *Drosophila* during exposure to thermal stress. Biol. J. Linn. Soc., 83, 197–205.

Fathy, R.F. (2024). Divergent perspectives on the synergistic impacts of thermal-chemical stress on aquatic biota within the framework of climate change scenarios. Chemosphere, 355, 141810.

Feldhaar, H. (2011). Bacterial symbionts as mediators of ecologically important traits of insect hosts. Ecol. Entomol., 36, 533–543.

Ferrer, A., Dorn, S. & Mazzi, D. (2013). Cross-generational effects of temperature on flight performance, and associated life-history traits in an insect. J. Evol. Biol., 26, 2321– 2330.

Fitzpatrick, J.L. & Lüpold, S. (2014). Sexual selection and the evolution of sperm quality. Mol. Hum. Reprod., 20, 1180–1189.

Folt, C., Chen, C., Moore, M. & Burnaford, J. (1999). Synergism and antagonism among multiple stressors. Limnol. Oceanogr., 44, 864–877.

Fournier-Level, A., Neumann-Mondlak, A., Good, R., Green, L., Schmidt, J.M. & Robin, C. (2016). Behavioural response to combined insecticide and temperature stress in natural populations of *Drosophila melanogaster*. J. Evol. Biol., 29, 1030–1044.

Frantzios, G., Paptsiki, K., Sidiropoulou, B., Lazaridis, I., Theophilidis, G. & Mavragani-Tsipidou, P. (2008). Evaluation of insecticidal and genotoxic effects of imidacloprid and acetochlor in *Drosophila melanogaster*. J. Appl. Entomol., 132, 583–590.

Freilich, S., Zarecki, R., Eilam, O., Segal, E.S., Henry, C.S., Kupiec, M., et al. (2011). Competitive and cooperative metabolic interactions in bacterial communities. Nat. Commun., 2, 589.

Gandara, A.C.P. & Drummond-Barbosa, D. (2022). Warm and cold temperatures have distinct germline stem cell lineage effects during *Drosophila oogenesis*. Development, 149, dev200149.

Gandara, A.C.P. & Drummond-Barbosa, D. (2023). Chronic exposure to warm temperature causes low sperm abundance and quality in *Drosophila melanogaster*. Sci. Rep., 13, 12331.

Gardner, J.L., Peters, A., Kearney, M.R., Joseph, L. & Heinsohn, R. (2011). Declining body size: a third universal response to warming? Trends Ecol. Evol., 26, 285–291.

Ge, L., Huang, L., Yang, G., Song, Q., Stanley, D., Gurr, G., et al. (2013). Molecular basis for insecticide-enhanced thermotolerance in the brown planthopper *Nilaparvata lugens Stål* (Hemiptera: Delphacidae). Mol. Ecol., 22, 5624–5634.

Gould, A.L., Zhang, V., Lamberti, L., Jones, E.W., Obadia, B., Korasidis, N., et al. (2018). Microbiome interactions shape host fitness. Proc. Natl. Acad. Sci., 115, E11951– E11960.

Gruntenko, N.Е., Ilinsky, Y.Yu., Adonyeva, N.V., Burdina, E.V., Bykov, R.A., Menshanov, P.N., et al. (2017). Various *Wolbachia* genotypes differently influence host *Drosophila* dopamine metabolism and survival under heat stress conditions. BMC Evol. Biol., 17, 252.

Gu, X., Tian, S., Wang, D., Fei, G. & Wei, H. (2010). Interaction between short-term heat pretreatment and fipronil on 2nd instar larvae of diamondback moth, *Plutella xylostella* (Linn). Dose-Response, 8, 331–346.

Guilhot, R., Xuéreb, A. & Fellous, S. (2025). Transmission of yeast and bacterial symbionts between sexual partners in *Drosophila suzukii* and *Drosophila melanogaster*. R. Soc. Open Sci., 12, 241149.

Guillaume, A.S., Monro, K. & Marshall, D.J. (2016). Transgenerational plasticity and environmental stress: do paternal effects act as a conduit or a buffer? Funct. Ecol., 30, 1175–1184.

Hallman, T.A. & Brooks, M.L. (2015). The deal with diel: Temperature fluctuations, asymmetrical warming, and ubiquitous metals contaminants. Environ. Pollut. Barking Essex 1987, 206, 88–94.

He, Y., Zhao, J., Wu, D., Wyckhuys, K.A. & Wu, K. (2011). Sublethal effects of imidacloprid on *Bemisia tabaci* (Hemiptera: Aleyrodidae) under laboratory conditions. J. Econ. Entomol., 104, 833–838.

Heino, J., Virkkala, R. & Toivonen, H. (2009). Climate change and freshwater biodiversity: detected patterns, future trends and adaptations in northern regions. Biol. Rev., 84, 39– 54.

Henry, Y., Overgaard, J. & Colinet, H. (2020). Dietary nutrient balance shapes phenotypic traits of *Drosophila melanogaster* in interaction with gut microbiota. Comp. Biochem. Physiol. A. Mol. Integr. Physiol., 241, 110626.

Hoekstra, L.A., Siddiq, M.A. & Montooth, K.L. (2013). Pleiotropic effects of a mitochondrial– nuclear incompatibility depend upon the accelerating effect of temperature in *Drosophila*. Genetics, 195, 1129–1139.

Holmstrup, M., Bindesbøl, A.-M., Oostingh, G.J., Duschl, A., Scheil, V., Köhler, H.-R., et al. (2010). Interactions between effects of environmental chemicals and natural stressors: a review. Sci. Total Environ., 408, 3746–3762.

Hooper, M.J., Ankley, G.T., Cristol, D.A., Maryoung, L.A., Noyes, P.D. & Pinkerton, K.E. (2013). Interactions between chemical and climate stressors: A role for mechanistic toxicology in assessing climate change risks. Environ. Toxicol. Chem., 32, 32–48.

Ihara, M., D. Buckingham, S., Matsuda, K. & B. Sattelle, D. (2017). Modes of action, resistance and toxicity of insecticides targeting nicotinic acetylcholine receptors. Curr. Med. Chem., 24, 2925–2934.

Iossa, G. (2019). Sex-specific differences in thermal fertility Limits. Trends Ecol. Evol., 34, 490–492.

Jaramillo, A. & Castañeda, L.E. (2021). Gut microbiota of *Drosophila subobscura* contributes to its heat tolerance and is sensitive to transient thermal stress. Front. Microbiol., 12, 654108.

Jones, R.M., Desai, C., Darby, T.M., Luo, L., Wolfarth, A.A., Scharer, C.D., et al. (2015). *Lactobacilli* modulate epithelial cytoprotection through the Nrf2 pathway. Cell Rep., 12, 1217–1225.

Jutfelt, F., Norin, T., Åsheim, E.R., Rowsey, L.E., Andreassen, A.H., Morgan, R., et al. (2021). ‘Aerobic scope protection’reduces ectotherm growth under warming. Funct. Ecol., 35, 1397–1407.

Kang, Z.-W., Liu, F.-H., Pang, R.-P., Tian, H.-G. & Liu, T.-X. (2018). Effect of sublethal doses of imidacloprid on the biological performance of aphid endoparasitoid *Aphidius gifuensis* (Hymenoptera: Aphidiidae) and influence on its related gene expression. Front. Physiol., 9, 1729.

Karimzadeh, R., Javanshir, M. & Hejazi, M.J. (2020). Individual and combined effects of insecticides, inert dusts and high temperatures on *Callosobruchus maculatus* (Coleoptera: Chrysomelidae). J. Stored Prod. Res., 89, 101693.

Kikuchi, Y., Hayatsu, M., Hosokawa, T., Nagayama, A., Tago, K. & Fukatsu, T. (2012). Symbiont-mediated insecticide resistance. Proc. Natl. Acad. Sci., 109, 8618–8622.

Kim, H.Y., Yu, S., Jeong, T. & Kim, S.D. (2014). Relationship between trans-generational effects of tetracycline on *Daphnia magna* at the physiological and whole organism level. Environ. Pollut., 191, 111–118.

Kimberly, D.A. & Salice, C.J. (2015). Multigenerational contaminant exposures produce non-monotonic, transgenerational responses in *Daphnia magna*. Environ. Pollut. Barking Essex 1987, 207, 176–182.

Kokou, F., Sasson, G., Nitzan, T., Doron-Faigenboim, A., Harpaz, S., Cnaani, A., et al. (2018). Host genetic selection for cold tolerance shapes microbiome composition and modulates its response to temperature. elife, 7, e36398.

Kooĳman, S.A.L.M. (2000). Dynamic energy and mass budgets in biological systems. Cambridge University Press.

Koyle, M.L., Veloz, M., Judd, A.M., Wong, A.C.-N., Newell, P.D., Douglas, A.E., et al. (2016). Rearing the fruit fly *Drosophila melanogaster* under axenic and gnotobiotic conditions. JoVE J. Vis. Exp. e54219.

Lenth, R.V. (2025). emmeans: Estimated marginal means, aka least-squares means. R package version 1.11.1, <https://CRAN.R-project.org/package=emmeans>.

Levy, R. & Borenstein, E. (2013). Metabolic modeling of species interaction in the human microbiome elucidates community-level assembly rules. Proc. Natl. Acad. Sci., 110, 12804–12809.

Li, S., Tan, H.-Y., Wang, N., Zhang, Z.-J., Lao, L., Wong, C.-W., et al. (2015). The role of oxidative stress and antioxidants in liver diseases. Int. J. Mol. Sci., 16, 26087–26124.

Li, W., Lu, Z., Li, L., Yu, Y., Dong, S., Men, X., et al. (2018). Sublethal effects of imidacloprid on the performance of the bird cherry-oat aphid *Rhopalosiphum padi*. PLOS ONE, 13, e0204097.

Liu, H., He, J., Chi, C. & Shao, J. (2014). Differential HSP70 expression in *Mytilus coruscus* under various stressors. Gene, 543, 166–173.

Lüdecke, D., Ben-Shachar, M., Patil, I., Waggoner, P. & Makowski, D. (2021). “performance: An R package for assessment, comparison and testing of statistical models.” J. Open Source Softw., 6, 3139.

Lüpold, S., Manier, M.K., Berben, K.S., Smith, K.J., Daley, B.D., Buckley, S.H., et al. (2012). How multivariate ejaculate traits determine competitive fertilization success in *Drosophila melanogaster*. Curr. Biol., 22, 1667–1672.

Lüpold, S., Reil, J.B., Manier, M.K., Zeender, V., Belote, J.M. & Pitnick, S. (2020). How female × male and male × male interactions influence competitive fertilization in *Drosophila melanogaster*. Evol. Lett., 4, 416–429.

Macaulay, S.J., Buchwalter, D.B. & Matthaei, C.D. (2020). Water temperature interacts with the insecticide imidacloprid to alter acute lethal and sublethal toxicity to mayfly larvae. N. Z. J. Mar. Freshw. Res., 54, 115–130.

Maggu, K., Meena, A., De Nardo, A.N., Sbilordo, S.H., Roy, J. & Lüpold, S. (2025). Male gut microbiome mediates postmating sexual selection in *Drosophila*. bioRxiv. doi: 10.1101/2025.05.23.655737.

Magnúsdóttir, S., Heinken, A., Kutt, L., Ravcheev, D.A., Bauer, E., Noronha, A., et al. (2017). Generation of genome-scale metabolic reconstructions for 773 members of the human gut microbiota. Nat. Biotechnol., 35, 81–89.

Manzi, C., Vergara-Amado, J., Franco, L.M. & Silva, A.X. (2020). The effect of temperature on candidate gene expression in the brain of honey bee *Apis mellifera* (Hymenoptera: Apidae) workers exposed to neonicotinoid imidacloprid. J. Therm. Biol., 93, 102696.

Mao, K., Jin, R., Li, W., Ren, Z., Qin, X., He, S., et al. (2019). The influence of temperature on the toxicity of insecticides to *Nilaparvata lugens* (Stål). Pestic. Biochem. Physiol., 156, 80–86.

Marshall, D.J., Heppell, S.S., Munch, S.B. & Warner, R.R. (2010). The relationship between maternal phenotype and offspring quality: do older mothers really produce the best offspring? Ecology, 91, 2862–2873.

Martelli, F., Zhongyuan, Z., Wang, J., Wong, C.-O., Karagas, N.E., Roessner, U., et al. (2020). Low doses of the neonicotinoid insecticide imidacloprid induce ROS triggering neurological and metabolic impairments in *Drosophila*. Proc. Natl. Acad. Sci., 117, 25840–25850.

Massot, M., Bagni, T., Maria, A., Couzi, P., Drozdz, T., Malbert-Colas, A., et al. (2021). Combined influences of transgenerational effects, temperature and insecticide on the moth *Spodoptera littoralis*. Environ. Pollut., 289, 117889.

Meena, A., De Nardo, A.N., Maggu, K., Sbilordo, S.H., Roy, J., Snook, R.R., et al. (2024a). Fertility loss and recovery dynamics after repeated heat stress across life stages in male *Drosophila melanogaster*: patterns and processes. R. Soc. Open Sci., 11, 241082.

Meena, A., Maggu, K., De Nardo, A.N., Sbilordo, S.H., Eggs, B., Al Toma Sho, R., et al. (2024b). Life stage-specific effects of heat stress on spermatogenesis and oogenesis in *Drosophila melanogaster*. J. Therm. Biol., 125, 104001.

Meng, S., Tran, T.T., Van Dinh, K., Delnat, V. & Stoks, R. (2022). Acute warming increases pesticide toxicity more than transgenerational warming by reducing the energy budget. Sci. Total Environ., 805, 150373.

Moeller, A.H., Ivey, K., Cornwall, M.B., Herr, K., Rede, J., Taylor, E.N., et al. (2020). The lizard gut microbiome changes with temperature and is associated with heat tolerance. Appl. Environ. Microbiol., 86, e01181–20.

Moghadam, N.N., Thorshauge, P.M., Kristensen, T.N., de Jonge, N., Bahrndorff, S., Kjeldal, H., et al. (2018). Strong responses of *Drosophila melanogaster* microbiota to developmental temperature. Fly (Austin), 12, 1–12.

Montllor, C.B., Maxmen, A. & Purcell, A.H. (2002). Facultative bacterial endosymbionts benefit pea aphids *Acyrthosiphon pisum* under heat stress. Ecol. Entomol., 27, 189–195.

Moore, M.P., Whiteman, H.H. & Martin, R.A. (2019). A mother’s legacy: the strength of maternal effects in animal populations. Ecol. Lett., 22, 1620–1628.

Moreira, D.R., de Souza, T.H.S., Galhardo, D., Puentes, S.M.D., Figueira, C.L., Silva, B.G. da, et al. (2022). Imidacloprid induces histopathological damage in the midgut, ovary, and spermathecal stored spermatozoa of queens after chronic colony exposure. Environ. Toxicol. Chem., 41, 1637–1648.

Morgan, H.L. & Watkins, A.J. (2019). Transgenerational impact of environmental change. In: Reproductive Sciences in Animal Conservation, Advances in Experimental Medicine and Biology (eds. Comizzoli, P., Brown, J.L. & Holt, W.V.). Springer International Publishing, Cham, pp. 71–89.

Mousseau, T.A., Uller, T., Wapstra, E. & Badyaev, A.V. (2009). Evolution of maternal effects: past and present. Philos. Trans. R. Soc. B Biol. Sci., 364, 1035–1038.

Navarro-Ortega, A., Acuña, V., Bellin, A., Burek, P., Cassiani, G., Choukr-Allah, R., et al. (2015). Managing the effects of multiple stressors on aquatic ecosystems under water scarcity. The GLOBAQUA project. Sci. Total Environ., 503–504, 3–9.

Neven, L.G. (2000). Physiological responses of insects to heat. Postharvest Biol. Technol., 21, 103–111.

Newell, P.D. & Douglas, A.E. (2014). Interspecies interactions determine the impact of the gut microbiota on nutrient allocation in *Drosophila melanogaster*. Appl. Environ. Microbiol., 80, 788–796.

Noecker, C., Chiu, H.-C., McNally, C.P. & Borenstein, E. (2019). Defining and evaluating microbial contributions to metabolite variation in microbiome-metabolome association studies. mSystems, 4, 10–1128.

Noyes, P.D. & Lema, S.C. (2015). Forecasting the impacts of chemical pollution and climate change interactions on the health of wildlife. Curr. Zool., 61, 669–689.

Noyes, P.D., McElwee, M.K., Miller, H.D., Clark, B.W., Van Tiem, L.A., Walcott, K.C., et al. (2009). The toxicology of climate change: environmental contaminants in a warming world. Environ. Int., 35, 971–986.

Obadia, B., Güvener, Z.T., Zhang, V., Ceja-Navarro, J.A., Brodie, E.L., William, W.J., et al. (2017). Probabilistic invasion underlies natural gut microbiome stability. Curr. Biol., 27, 1999–2006.

Oksanen, J., Simpson, G., Blanchet, F., Kindt, R., Legendre, P., Minchin, P., et al. (2025). vegan: Community ecology package. R package version 2.7–1.

Osborne, B., Siboni, N., Seymour, J.R., Ralph, P. & Pernice, M. (2023). Exploring the potential of algae-bacteria interactions in the biocontrol of the marine pathogen *Vibrio parahaemolyticus*. J. Appl. Phycol., 35, 2731–2743.

Overgaard, J., Malmendal, A., Sørensen, J.G., Bundy, J.G., Loeschcke, V., Nielsen, N.C., et al. (2007). Metabolomic profiling of rapid cold hardening and cold shock in *Drosophila melanogaster*. J. Insect Physiol., 53, 1218–1232.

Paaĳmans, K.P., Heinig, R.L., Seliga, R.A., Blanford, J.I., Blanford, S., Murdock, C.C., et al. (2013). Temperature variation makes ectotherms more sensitive to climate change. Glob. Change Biol., 19, 2373–2380.

Pang, R., Chen, M., Yue, L., Xing, K., Li, T., Kang, K., et al. (2018). A distinct strain of Arsenophonus symbiont decreases insecticide resistance in its insect host. PLoS Genet., 14, e1007725.

Parker, G.A. & Pizzari, T. (2010). Sperm competition and ejaculate economics. Biol. Rev., 85, 897–934.

Piggott, J.J., Niyogi, D.K., Townsend, C.R. & Matthaei, C.D. (2015a). Multiple stressors and stream ecosystem functioning: climate warming and agricultural stressors interact to affect processing of organic matter. J. Appl. Ecol., 52, 1126–1134.

Piggott, J.J., Townsend, C.R. & Matthaei, C.D. (2015b). Climate warming and agricultural stressors interact to determine stream macroinvertebrate community dynamics. Glob. Change Biol., 21, 1887–1906.

Pilakouta, N. & Baillet, A. (2022). Effects of temperature on mating behaviour and mating success: A meta-analysis. J. Anim. Ecol., 91, 1642–1650.

Pizzari, T. & Parker, G.A. (2009). Sperm competition and sperm phenotype. In: Sperm Biology. Elsevier, pp. 207–245.

Pölkki, M., Kangassalo, K. & Rantala, M.J. (2012). Transgenerational effects of heavy metal pollution on immune defense of the blow fly *Protophormia terraenovae*. PLoS One, 7, e38832.

Pörtner, H.-O., Bock, C. & Mark, F.C. (2017). Oxygen-and capacity-limited thermal tolerance: bridging ecology and physiology. J. Exp. Biol., 220, 2685–2696.

Przeslawski, R., Byrne, M. & Mellin, C. (2015). A review and meta-analysis of the effects of multiple abiotic stressors on marine embryos and larvae. Glob. Change Biol., 21, 2122– 2140.

Reátegui-Zirena, E.G., Fidder, B.N., Olson, A.D., Dawson, D.E., Bilbo, T.R. & Salice, C.J. (2017). Transgenerational endpoints provide increased sensitivity and insight into multigenerational responses of *Lymnaea stagnalis* exposed to cadmium. Environ. Pollut., 224, 572–580.

Renoz, F., Pons, I. & Hance, T. (2019). Evolutionary responses of mutualistic insect–bacterial symbioses in a world of fluctuating temperatures. Curr. Opin. Insect Sci., 35, 20–26.

Ricupero, M., Abbes, K., Haddi, K., Kurtulus, A., Desneux, N., Russo, A., et al. (2020). Combined thermal and insecticidal stresses on the generalist predator *Macrolophus pygmaeus*. Sci. Total Environ., 729, 138922.

Ridley, E.V., Wong, A.C. & Douglas, A.E. (2013). Microbe-dependent and nonspecific effects of procedures to eliminate the resident microbiota from *Drosophila melanogaster*. Appl. Environ. Microbiol., 79, 3209–3214.

Ridley, E.V., Wong, A.C., Westmiller, S. & Douglas, A.E. (2012). Impact of the resident microbiota on the nutritional phenotype of *Drosophila melanogaster*. PloS One, 7, e36765.

Roy, P., Phukan, P.K., Changmai, D. & Boruah, S. (2017). Pesticides, insecticides and male infertility. Int J Reprod Contracept Obs Gynecol, 6, 3387–3391.

Russell, J.A. & Moran, N.A. (2006). Costs and benefits of symbiont infection in aphids: variation among symbionts and across temperatures. Proc. Biol. Sci., 273, 603–610.

Sales, K., Thomas, P., Gage, M.J. & Vasudeva, R. (2024). Experimental heatwaves reduce the effectiveness of ejaculates at occupying female reproductive tracts in a model insect. R. Soc. Open Sci., 11, 231949.

Sales, K., Vasudeva, R., Dickinson, M.E., Godwin, J.L., Lumley, A.J., Michalczyk, Ł., et al. (2018). Experimental heatwaves compromise sperm function and cause transgenerational damage in a model insect. Nat. Commun., 9, 4771.

Schultz, C.L., Wamucho, A., Tsyusko, O.V., Unrine, J.M., Crossley, A., Svendsen, C., et al. (2016). Multigenerational exposure to silver ions and silver nanoparticles reveals heightened sensitivity and epigenetic memory in *Caenorhabditis elegans*. Proc. R. Soc. B Biol. Sci., 283, 20152911.

Schwasinger-schmidt, T.E., Kachman, S.D. & Harshman, L.G. (2012). Evolution of starvation resistance in *Drosophila melanogaster*: measurement of direct and correlated responses to artificial selection. J. Evol. Biol., 25, 378–387.

Segner, H., Schmitt-Jansen, M. & Sabater, S. (2014). Assessing the impact of multiple stressors on aquatic biota: The receptor’s side matters. Environ. Sci. Technol., 48, 7690–7696.

Semsar-Kazerouni, M., Boerrigter, J.G. & Verberk, W.C. (2020). Changes in heat stress tolerance in a freshwater amphipod following starvation: The role of oxygen availability, metabolic rate, heat shock proteins and energy reserves. Comp. Biochem. Physiol. A. Mol. Integr. Physiol., 245, 110697.

Sepulveda, J. & Moeller, A.H. (2020). The effects of temperature on animal gut microbiomes. Front. Microbiol., 11, 384.

Shahid, N., Siddique, A. & Liess, M. (2024). Synergistic interaction between a toxicant and food stress is further exacerbated by temperature. Environ. Pollut., 363, 125109.

Shama, L.N., Strobel, A., Mark, F.C. & Wegner, K.M. (2014). Transgenerational plasticity in marine sticklebacks: maternal effects mediate impacts of a warming ocean. Funct. Ecol., 28, 1482–1493.

Shokal, U., Yadav, S., Atri, J., Accetta, J., Kenney, E., Banks, K., et al. (2016). Effects of co-occurring *Wolbachia* and *Spiroplasma* endosymbionts on the *Drosophila* immune response against insect pathogenic and non-pathogenic bacteria. BMC Microbiol., 16, 16.

Silva, A.P.N., Andrade, E.S., Nascimento, V.L. & Haddi, K. (2025). Thermal modulation of insecticide-induced hormetic and oxidative responses in insect pests. Chemosphere, 370, 143920.

Silva, A.S.J., Kristiansen, S.M., Sengupta, S., van Gestel, C.A., Leinaas, H.P. & Borgå, K. (2023). Using dietary exposure to determine sub-lethal effects from imidacloprid in two springtail (*Collembola*) species. Ecotoxicology, 32, 1209–1220.

Simmons, L.W. & Fitzpatrick, J.L. (2012). Sperm wars and the evolution of male fertility. Reproduction, 144, 519–534.

Siviter, H., Bailes, E.J., Martin, C.D., Oliver, T.R., Koricheva, J., Leadbeater, E., et al. (2021). Agrochemicals interact synergistically to increase bee mortality. Nature, 596, 389–392.

Sokolova, I.M. (2013). Energy-limited tolerance to stress as a conceptual framework to integrate the effects of multiple stressors. Integr. Comp. Biol., 53, 597–608.

Sørensen, J.G., Kristensen, T.N. & Loeschcke, V. (2003). The evolutionary and ecological role of heat shock proteins. Ecol. Lett., 6, 1025–1037.

Stehle, S. & Schulz, R. (2015). Agricultural insecticides threaten surface waters at the global scale. Proc. Natl. Acad. Sci., 112, 5750–5755.

Tran, T.T., Janssens, L., Dinh, K.V. & Stoks, R. (2018). Transgenerational interactions between pesticide exposure and warming in a vector mosquito. Evol. Appl., 11, 906– 917.

Uller, T., Nakagawa, S. & English, S. (2013). Weak evidence for anticipatory parental effects in plants and animals. J. Evol. Biol., 26, 2161–2170.

Verberk, W.C.E.P., Overgaard, J., Ern, R., Bayley, M., Wang, T., Boardman, L., et al. (2016). Does oxygen limit thermal tolerance in arthropods? A critical review of current evidence. Comp. Biochem. Physiol. A. Mol. Integr. Physiol., 192, 64–78.

Verheyen, J. & Stoks, R. (2019). Current and future daily temperature fluctuations make a pesticide more toxic: Contrasting effects on life history and physiology. Environ. Pollut., 248, 209–218.

Verheyen, J. & Stoks, R. (2023). Thermal performance curves in a polluted world: too cold and too hot temperatures synergistically increase pesticide toxicity. Environ. Sci. Technol., 57, 3270–3279.

Walsh, B.S., Parratt, S.R., Atkinson, D., Snook, R.R., Bretman, A. & Price, T.A. (2019). Integrated approaches to studying male and female thermal fertility limits. Trends Ecol. Evol., 34, 492–493.

Walsh, M.R., Whittington, D. & Funkhouser, C. (2014). Thermal transgenerational plasticity in natural populations of *Daphnia*.

Wan, N.-F., Fu, L., Dainese, M., Kiær, L.P., Hu, Y.-Q., Xin, F., et al. (2025). Pesticides have negative effects on non-target organisms. Nat. Commun., 16, 1360.

Wang, X., Ma, D., Jin, Q., Deng, S., Stančin, H., Tan, H., et al. (2019). Synergistic effects of biomass and polyurethane co-pyrolysis on the yield, reactivity, and heating value of biochar at high temperatures. Fuel Process. Technol., 194, 106127.

Wang, Y.-C., Chang, Y.-W., Bai, J., Zhang, X.-X., Iqbal, J., Lu, M.-X., et al. (2021a). High temperature stress induces expression of CYP450 genes and contributes to insecticide tolerance in *Liriomyza trifolii*. Pestic. Biochem. Physiol., 174, 104826.

Wang, Y.-C., Chang, Y.-W., Bai, J., Zhang, X.-X., Iqbal, J., Lu, M.-X., et al. (2021b). Temperature affects the tolerance of *Liriomyza trifolii* to insecticide abamectin. Ecotoxicol. Environ. Saf., 218, 112307.

Wickham, H. (2016). *ggplot2: Elegant Graphics for Data Analysis*. Springer, New York.

Xiao, D., Zhao, J., Guo, X., Chen, H., Qu, M., Zhai, W., et al. (2016). Sublethal effects of imidacloprid on the predatory seven-spot ladybird beetle *Coccinella septempunctata*. Ecotoxicology, 25, 1782–1793.

Yu, C.-W. & Liao, V.H.-C. (2016). Transgenerational Reproductive effects of arsenite are associated with H3K4 dimethylation and SPR-5 downregulation in *Caenorhabditis elegans*. Environ. Sci. Technol., 50, 10673–10681.

Zelezniak, A., Andrejev, S., Ponomarova, O., Mende, D.R., Bork, P. & Patil, K.R. (2015). Metabolic dependencies drive species co-occurrence in diverse microbial communities. Proc. Natl. Acad. Sci., 112, 6449–6454.

Zhang, X., Yu, X. & Chen, J. (2008). High temperature effects on yeast-like endosymbiotes and pesticide resistance of the small brown planthopper, *Laodelphax striatellus*. Rice Sci., 15, 326–330.

Zhang, Y., Cai, T., Ren, Z., Liu, Y., Yuan, M., Cai, Y., et al. (2021a). Decline in symbiont-dependent host detoxification metabolism contributes to increased insecticide susceptibility of insects under high temperature. ISME J., 15, 3693–3703.

Zhang, Y., Xu, G., Jiang, Y., Ma, C. & Yang, G. (2021b). Sublethal effects of Imidacloprid on fecundity, apoptosis and virus transmission in the small brown planthopper *Laodelphax striatellus*. Insects, 12, 1131.

Zhou, C., Liu, L., Yang, H., Wang, Z., Long, G. & Jin, D. (2017). Sublethal effects of imidacloprid on the development, reproduction, and susceptibility of the white-backed planthopper, Sogatella furcifera (Hemiptera: Delphacidae). J. Asia-Pac. Entomol., 20, 996–1000.

Zwoinska, M.K., Rodrigues, L.R., Slate, J. & Snook, R.R. (2020). Phenotypic responses to and genetic architecture of sterility following exposure to sub-lethal temperature during development. Front. Genet., 11, 573.

